# Immune History Modifies Disease Severity to HPAI H5N1 Clade 2.3.4.4b Viral Challenge

**DOI:** 10.1101/2024.10.23.619695

**Authors:** Pamela H. Brigleb, Bridgett Sharp, Ericka Kirkpatrick Roubidoux, Victoria Meliopoulos, Shaoyuan Tan, Brandi Livingston, Dorothea Morris, Tyler Ripperger, Lauren Lazure, Velmurugan Balaraman, Alexis C. Thompson, Katie Kleinhenz, Kiril Dimitrov, Paul G. Thomas, Stacey Schultz-Cherry

## Abstract

The most recent outbreak of highly pathogenic avian H5 influenza (HPAI) virus in cattle is now widespread across the U.S. with spillover events happening to other mammals, including humans. Several human cases have been reported with clinical signs ranging from conjunctivitis to respiratory illness. However, most of those infected report mild to moderate symptoms, while previously reported HPAI H5Nx infections in humans have had mortality rates upwards of 50%. We recently reported that mice with pre-existing immunity to A/Puerto Rico/08/1934 H1N1 virus were protected from lethal challenge from highly pathogenic clade 2.3.4.4b H5N1 influenza virus. Here, we demonstrate that mice infected with the 2009 pandemic H1N1 virus strain A/California/04/2009 (Cal09) or vaccinated with a live-attenuated influenza vaccine (LAIV) were moderately-to-highly protected against a lethal A/bovine/Ohio/B24OSU-439/2024 H5N1 virus challenge. We also observed that ferrets with mixed pre-existing immunity—either from LAIV vaccination and/or from Cal09 infection—showed protection against a HPAI H5N1 clade 2.3.4.4b virus isolated from a cat. Notably, this protection occurred independently of any detectable hemagglutination inhibition titers (HAIs) against the H5N1 virus. To explore factors that may contribute to protection, we conducted detailed T cell epitope mapping using previously published sequences from H1N1 strains. This analysis revealed a high conservation of amino acid sequences within the internal proteins of our bovine HPAI H5N1 virus strain. These data highlight the necessity to explore additional factors that contribute to protection against HPAI H5N1 viruses, such as memory T cell responses, in addition to HA-inhibition or neutralizing antibodies.

## Introduction

In March 2024, HPAI H5N1 virus clade 2.3.4.4b was discovered in dairy cows in the United States with spillover events to mammals including cats and humans (*1–3*). By September 20, 2024, this strain had been identified in 14 states, as reported by the American Veterinary Medical Association (*4*). There have been at least 14 confirmed human cases primarily in farm workers since the outbreak, with patients presenting with generally mild to moderate clinical symptoms (*5–7*). Previous spillover events with HPAI H5Nx viruses in humans have an average fatality rate of around 50% (*8*), and with the increasing case reports in humans and multiple avenues of crossover events to other mammals (*2*), it is important to understand mechanisms of protection against HPAI H5N1 clade 2.3.4.4b viruses.

A recent study in preprint found that persons born after the 1960’s do not contain neutralizing antibody responses to bovine HPAI H5N1 virus (*9*). This corroborates previous findings in 2016 that the elderly may have higher protection against H5N1 viruses, as they have been exposed to viral strains that better match previous H5N1 viruses (*10*). In ferrets, prior H1N1 virus infection immunity with a 2009 pandemic strain provided protection against lethal challenge of bovine HPAI H5N1 clade 2.3.4.4b virus independent of H5N1 specific neutralizing antibodies (*9*). This is corroborated by our previous findings in mice that prior infection with the H1N1 virus A/Puerto Rico/08/34 was sufficient for protection, also independent of H5N1 hemagglutinin (HA) specific neutralizing antibodies (*11*). These results indicate there are factors that contribute to protection against HPAI H5N1 viral infection independent of neutralizing antibodies, such as non-neutralizing antibodies and memory T cell responses, at least with older H1N1 influenza strains

In this study, we tested whether mice previously infected with the pandemic H1N1 A/California/04/2009 (Cal09) virus or vaccinated with the 2023-24 FluMist vaccine had protective immunity against HPAI bovine H5N1 clade 2.3.4.4b virus. We further tested whether ferrets with mixed pre-existing immunity – vaccination with an LAIV and/or H1N1 infection - were protected against lethal challenge of a HPAI H5N1 clade 2.3.4.4b virus isolated from a cat, which may have a greater spillover potential than the bovine-derivative. Our results show pre-existing immunity in both mice and ferrets provides some level of protection against lethal challenge of HPAI H5N1 clade 2.3.4.4b. While HAI responses were negative, non-neutralizing antibodies to whole virus were induced in mice and ferrets protected against HPAI H5N1 viruses. Further, in-depth analysis of known T cell epitopes shows conservation between H1N1 and the bovine H5N1 amino acid sequences suggesting that non-neutralizing antibodies and T cells may be important for protection from severe disease. Collectively, these studies highlight the necessity to explore additional factors that may contribute to protection against HPAI H5N1 viruses in addition to HA-inhibition or neutralizing antibodies.

## Results

### Prior H1N1 virus infection history or vaccination with LAIV provides mild to high protection against lethal bovine HPAI H5N1

We have previously demonstrated that prior infection with the H1N1 A/Puerto Rico/08/34 virus protected mice from lethal HPAI clade 2.3.4.4b bovine H5N1 infection (*11*). We were curious whether a more contemporary H1N1 strain, Cal09 virus, or a live-attenuated vaccine from the previous influenza season (FluMist, 2023-24) would also provide protection against lethal bovine HPAI H5N1 viral challenge. To test, a combination of adult male and female C57BL/6J mice were intranasally (IN) inoculated with 100 50% tissue culture infectious dose (TCID_50_) of Cal09 or ∼10^5^ particles of LAIV under light anesthesia and were monitored daily. At 3 weeks post-infection, pre-challenge sera was collected, and mice were challenged with 5x their respective mean lethal-dose

50 (mLD_50_) (*11*) of the bovine HPAI H5N1 virus to A/bovine/Ohio/B24OSU-439/2024 (A/bovine/Ohio/24) virus. The mice were split into three groups: two for tissue collection for viral titers on days 2 and 5 post-infection, and one for weight loss, clinical scores, and survival (**Figure 1A**). Pre-challenge sera was assessed for both hemagglutination inhibition (HAI) titers to H1N1 viruses (Cal09 and A/Victoria/2019 for LAIV) and for H5N1 antibodies by whole virus ELISA. This was selected since we and others have previously reported poor HAI detection and responses against bovine H5N1 infection (*11*). Most mice had either a positive H1N1 pre-challenge HAI titer (HAI titer ≥ 1:20) or a positive antibody response to H5N1 whole virus (≥ 1:100), and those that did not were excluded from the study (**Figure 1B, Supplemental Figure 1**).

**Figure 1.**
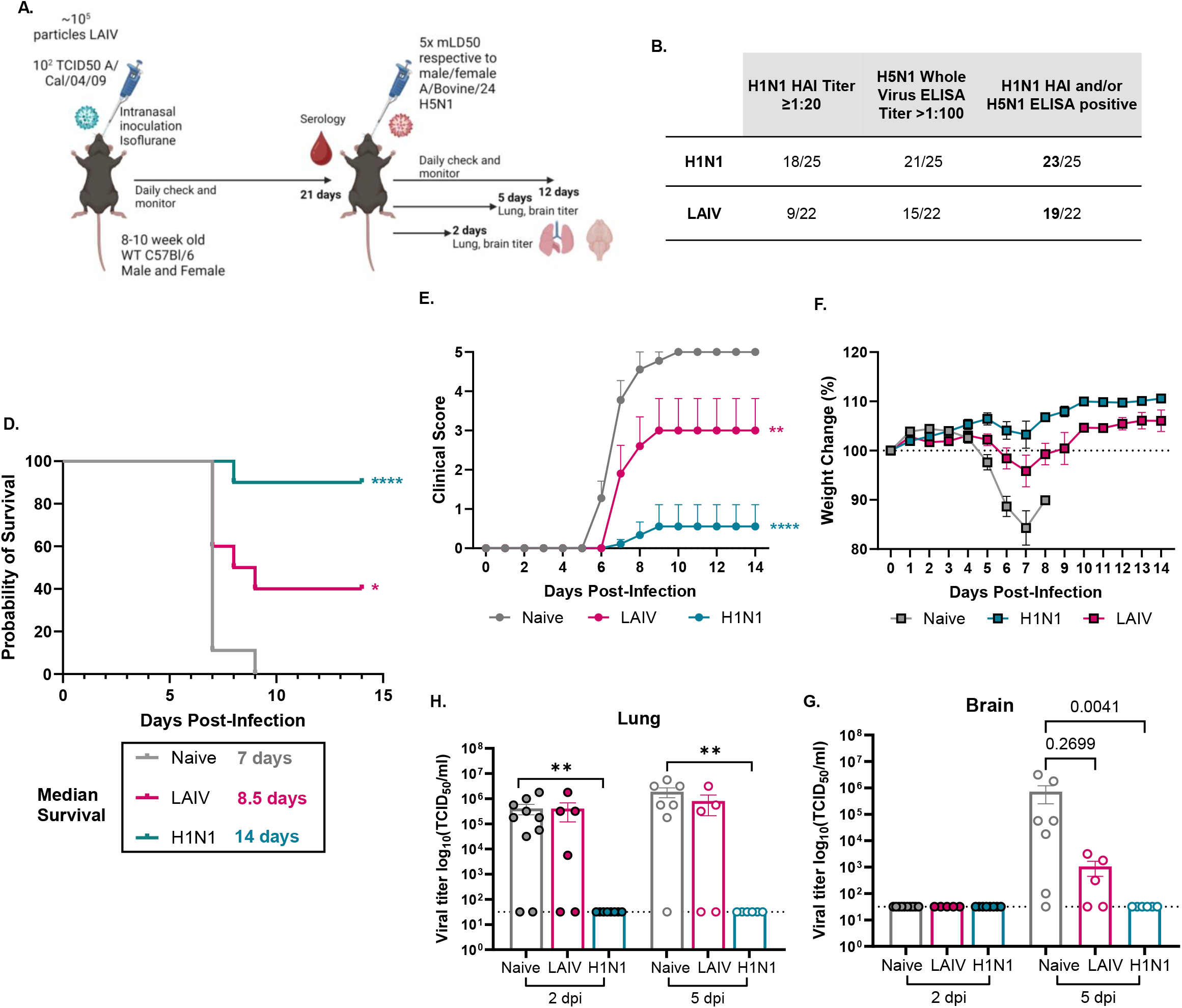
H1N1 prior infection and LAIV provides moderate to high protection against lethal bovine H5N1 viral challenge, respectively. **(A)** Graphical summary of experimental design made in *Biorender*. **(B)** Summary of pre-challenge serology against H1N1 or H5N1 viruses in immune history mice. **(C-E)** (n=9-10, 1 independent repeat) Mice were infected with 5x respective mLD_50_s (male and female) of A/bovine/Ohio/24 by IN inoculation and were monitored for survival **(C)**, clinical scores **(D)** and weight loss **(E)** or **(F, G)** mice were sacrificed at 2 or 5 dpi and lung **(F)** and brain **(G)** tissues were assessed for viral titers by TCID_50_ (n=5-10, 1 independent repeat). Statistical analyses include Log-rank Mantel Cox test (C), Two-way ANOVA (D, E) with Kruskal-Wallis Tests to account for repeated measurements (F, G) * p<.05; ** p < .01; **** p < .0001.

We found that prior infection with the H1N1 Cal09 virus or vaccination with LAIV provided significantly high to moderate protection against mortality from lethal bovine HPAI H5N1 virus, with 90% protection with H1N1 virus immune history and 40% protection with LAIV, compared to 0% survival in naïve mice (**Figure 1C**). We also observed protection against morbidity, with significantly decreased clinical scores (**Figure 1D**) and decreased weight loss (**Figure 1E**) compared to the naïve group. Strikingly, there were no detectable viral titers in the lung and brain at days 2 and 5 post-infection in the H1N1 virus immune history group (**Figures 1F and 1G**). While there were still detectable viral titers in the LAIV immune history group (**Figures 1F and 1G**), we detected a reduction in brain viral titer at 5 days post-infection (dpi) compared to the naïve group (**Figure 1G**). Overall, immune history from a H1N1 virus infection or LAIV vaccination provided some level of protection against mortality and morbidity from lethal bovine H5N1 viral challenge.

### Mixed Pre-Existing Immunity Provides Protection Against Lethal Challenge of a Feline-Derived HPAI H5N1 clade 2.3.4.4b virus

We isolated a clade 2.3.4.4b HPAI H5N1 from a cat found deceased near a dairy farm in New Mexico, A/feline/NewMexico/C001/2024 (A/feline/NewMexico/24). This strain is similar to but not identical to the A/bovine/Ohio/24 virus used in the murine immune history studies (**Table 1**). Since no studies to date have investigated whether immune history in ferrets also provides protection against feline derived HPAI H5N1 viruses, we IN inoculated naïve or immune history ferrets with 10^4^ TCID_50_ of A/feline/NewMexico/24 H5N1 virus isolated from a H5N1-infected cat and monitored them daily. One day-post infection, immune history direct contacts were cohoused with the index ferrets and were monitored daily (**Figure 2A**). The ferrets with immune history had been vaccinated with the 2010-11 season FluMist or an H1N1/H3N2 LAIV with a PR8 backbone currently being developed at St. Jude Children’s Research Hospital ∼6 months previously followed by challenge with PBS or A/Cal/09 H1N1 virus. See methods for a more detailed timeline.

**Table 1.** Amino acid sequence similarity amongst influenza A strains compared to A/bovine/Ohio/B24OSU-439/2024 including A/feline/NewMexico/0002/2024, A/Puerto Rico/8/1934, A/California/04/2009, A/Victoria/4897/2022, A/Wisconsin/67/2022, A/Ann Arbor/6/60, A/Norway/31694/2022.

|  | Bovine vs.<br>Feline H5N1 | Bovine vs. PR8 | Bovine vs.<br>Cal09 | Bovine vs.<br>Victoria | Bovine vs.<br>Wisconsin | Bovine vs.<br>Ann Arbor | Bovine vs.<br>Norway |
| --- | --- | --- | --- | --- | --- | --- | --- |
| **PB2** | 99.87% | 95.52% | 96.57% | 95.65% | 95.65% | 94.99% | 95.52% |
| **PB1** | 99.87% | 96.57% | 96.96% | 96.70% | 96.70% | 98.68% | 96.70% |
| **PA** | 99.72% | 95.81% | 96.37% | 95.53% | 95.67% | 94.97% | 95.67% |
| **NP** | 100% | 93.98% | 93.98% | 93.78% | 93.98% | 93.37% | 93.98% |
| **HA** | 100% | 64.22% | 62.61% | 61.78% | 62.13% | 71.55% | 61.95% |
| **NA** | 100% | 83.58% | 89.77% | 86.35% | 86.14% | 45.40% | 86.14% |
| **HA** | 100% | 64.22% | 62.61% | 61.78% | 62.13% | 71.55% | 61.95% |
| **M1** | 100% | 92.46% | 92.86% | 91.27% | 91.27% | 93.25% | 91.67% |
| **M2** | 100% | 83.51% | 88.66% | 86.60% | 86.60% | 81.44% | 86.60% |
| **NS1** | 99.13% | 91.30% | 79.13% | 76.09% | 76.52% | 85.65% | 76.52% |
| **Average Percent Similarity (including HA)** | 99.82% | 88.55% | 88.55% | 87.08% | 87.18% | 84.37% | 87.19% |
| **Average Percent Similarity (no HA)** | 99.82% | 91.59% | 91.79% | 90.25% | 90.32% | 85.97% | 90.35% |

**Figure 2.**
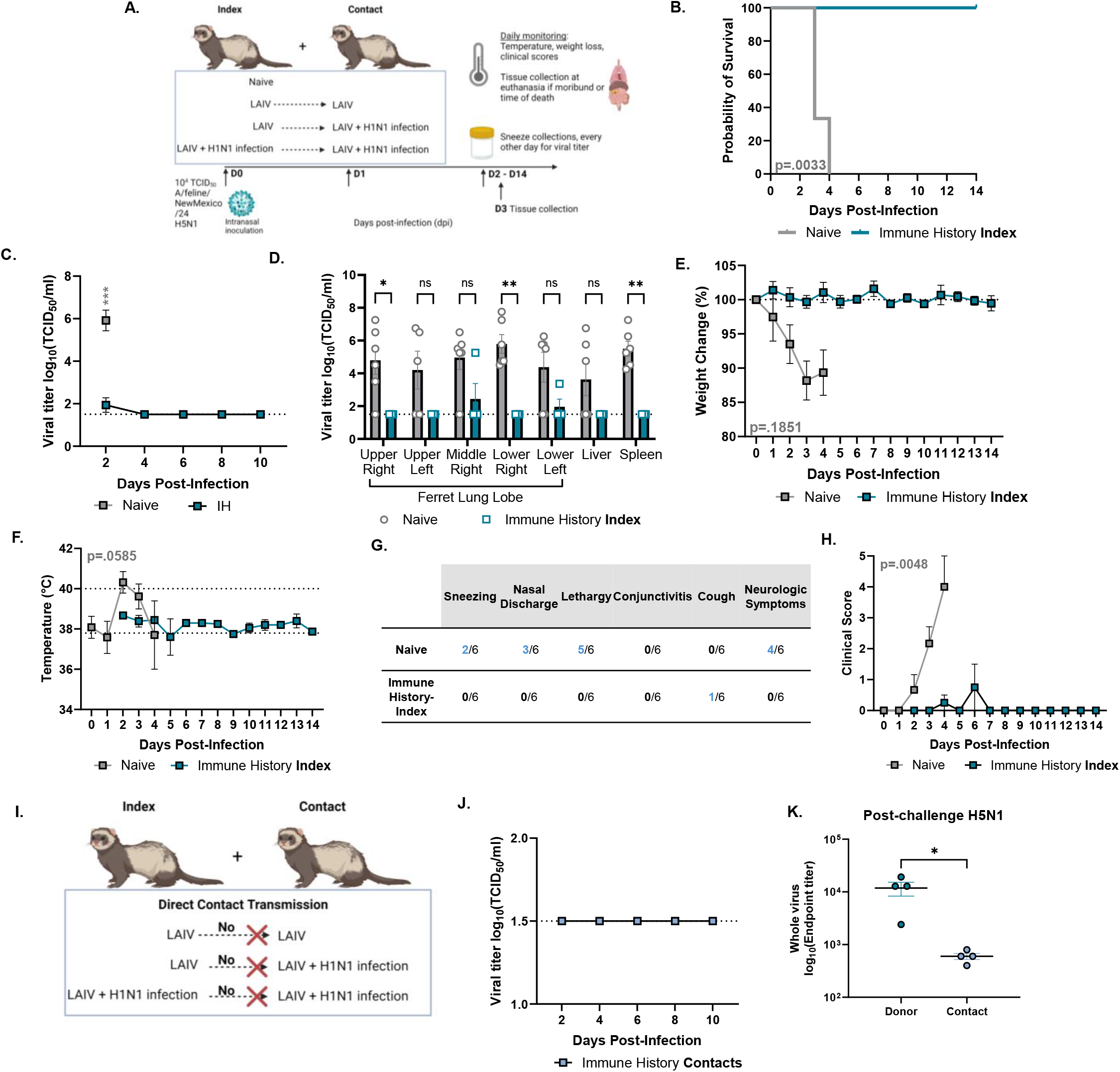
Mixed immunity provides protection against lethal challenge with feline-derived HPAI H5N1 A/Feline/NewMexico/24 strains in ferrets. **(A)** Graphical summary of experimental design made in *Biorender*. Ferrets were IN inoculated with 10^4^ TCID_50_ of A/Feline/NewMexico/24 and monitored daily (n=4-6 ferrets/group). **(B)** Survival following infection **(C)** tissues were harvested for viral titer by determination by TCID_50_ at 3-4 dpi **(D)** sneeze titers determined by TCID_50_ **(E)** weight loss **(F)** temperature **(G)** clinical scores and **(H)** summation of total clinical score daily. **(I)** Graphical summary of transmission experimental design made in *Biorender*, n=4 ferrets/group. **(J)** sneeze titers post co-housing in immune history contacts determined by TCID_50_ **(K)** sera antibody responses to bovine HPAI H5N1 by whole virus-ELISA 21 days post-H5N1 viral challenge and 20 days-post co-housing contacts and index ferrets.

A/feline/NewMexico/24 H5N1 virus caused 100% mortality in naïve ferrets by day 4 post-infection (**Figure 2B**). This was associated with high viral load in the upper respiratory tract and lungs with systemic spread **(Figure 2C**), significant clinical symptoms including weight loss, fever, cough (**Figures 2E-H**), and 4/6 ferrets exhibiting neurological symptoms prior to succumbing to infection or being humanely euthanized (**Figure 2G**). In contrast, ferrets with immune history were protected from HPAI A/feline/NewMexico/24 H5N1 viral challenge (**Figure 2B**). Little-to-no virus could be detected in sneezes with reduced systemic and lung viral titers (**Figure 2C and D**), minimal clinical signs with only one ferret out of six having a clinical symptom of a cough (**Figures 2E-H**) and no neurological disease (**Figure 2G**).

Previous studies have suggested that clade 2.3.4.4b HPAI H5N1 viruses do not efficiently transmit in ferret models. We also observed no effective transmission of the A/feline/NewMexico/24 H5N1 virus in immune history ferrets (**Figure 2I**) with no detectable virus in the sneezes of the immune history contacts at any time (**Figure 2J**). Furthermore, the index ferrets had significantly higher antibodies to whole H5N1 virus (A/bovine/Ohio/24) compared to the contact ferrets, indicating H5N1 infectivity in the index but not contact ferrets (**Figure 2K**). The bovine strain was used to assess serology as the assay was already optimized and the feline and bovine strains are 100% identical for the HA and NA proteins. The detection of positive antibody titers to H5N1 whole virus in the contacts is likely due to the cross-protective nature of the pre-existing immune antibodies as we determined in the mouse studies (**Figure 1**). Collectively, these findings indicate that immune history, from LAIV or a combination of LAIV and H1N1 infection, was sufficient to protect ferrets against lethal challenge of HPAI H5N1 virus isolated from a cat.

### Identifying Factors that Contribute to Protection Against HPAI H5N1 virus

We have shown that prior infection with H1N1 and LAIV vaccination offer protection against lethal HPAI H5N1 clade 2.3.4.4b challenges. However, neutralizing antibodies do not appear to correlate with this protection. Here, we demonstrate that immune-history mice exhibit cross-reactive antibodies to HPAI H5N1 prior to challenge, as measured by whole-virus ELISA (**Figure 1, Supplementary Figure 2**). These antibodies are primarily cross-reactive to the influenza surface proteins, hemagglutinin (HA) and neuraminidase (NA).

We compared the amino acid sequences of viral proteins from the feline H5N1 virus and several influenza A viruses represented in our immune-history studies - Cal09, PR8, and A/Ann Arbor/6/60, and A/Norway/31694/2022 (FluMist) – or viruses included in the seasonal influenza vaccine: A/Victoria/4897/2022 and A/Wisconsin/67/2022 (Table 1). Since the feline H5N1 isolate was highly similar to our bovine H5N1 isolate, exhibiting 99.82% sequence homology, we conducted the remainder of our studies with the bovine H5N1 virus strain.

Unsurprisingly, the most variable viral protein compared to the bovine H5N1 virus was HA, with the closest similarity being 71.55% to the H2N2 A/Ann Arbor/6/60 virus (**Table 1**). However, there is moderate conservation of the NA protein across HXN1 strains, with the highest similarity of 89.77% being to the Cal09 H1N1 virus (**Table 1**). Therefore, it is highly probable that the cross-protection observed in our immune-history mice against whole-virus ELISAs is attributable to the NA protein.

Upon conducting amino acid sequence alignments with other viral proteins, we noted a high conservation of internal proteins, including nucleoprotein (NP) and matrix protein 1 (M1), as well as RNA polymerase proteins such as PB1 and PB2 (**Table 1**). Several key T cell epitopes critical for protective responses against H1N1 viruses are known to reside within NP, PB1, and M1 (*12–15*). T cell memory responses to influenza are crucial for protection, alongside antibody responses. Given the high conservation of viral proteins containing well-characterized T cell epitopes, we next explored the potential for eliciting protective T cell immunity in our immune-history groups against HPAI H5N1 viral challenge.

### High Conversation of Previously Mapped T Cell Epitopes for Influenza A viruses

To assess whether previously mapped T cell epitopes from other influenza A viruses can provide cross-protection against HPAI H5N1 virus, we utilized the Immune Epitope Database & Tools interface (**Figure 3A**). We identified a total of 350 T cell epitopes for PR8 virus, 90 epitopes for Cal09 virus, and one for AnnArbor/60 virus, which we then separated based on mouse, human class I, or human class II restricted known epitopes. We then employed the Epitope Conservancy Analysis from the Immune Epitope Database to determine whether each known T cell epitope amino acid sequence was conserved in the bovine HPAI H5N1 virus. We categorized our results based on the level of amino acid conservation: 100% or ≥90% (averaging one amino acid dissimilarity). Notably, the NS1 protein human and mouse T cell epitope mapped for AnnArbor/60, the backbone of FluMist LAIV, was conserved across the bovine H5N1 strains (**Figure 3B**).

**Figure 3.**
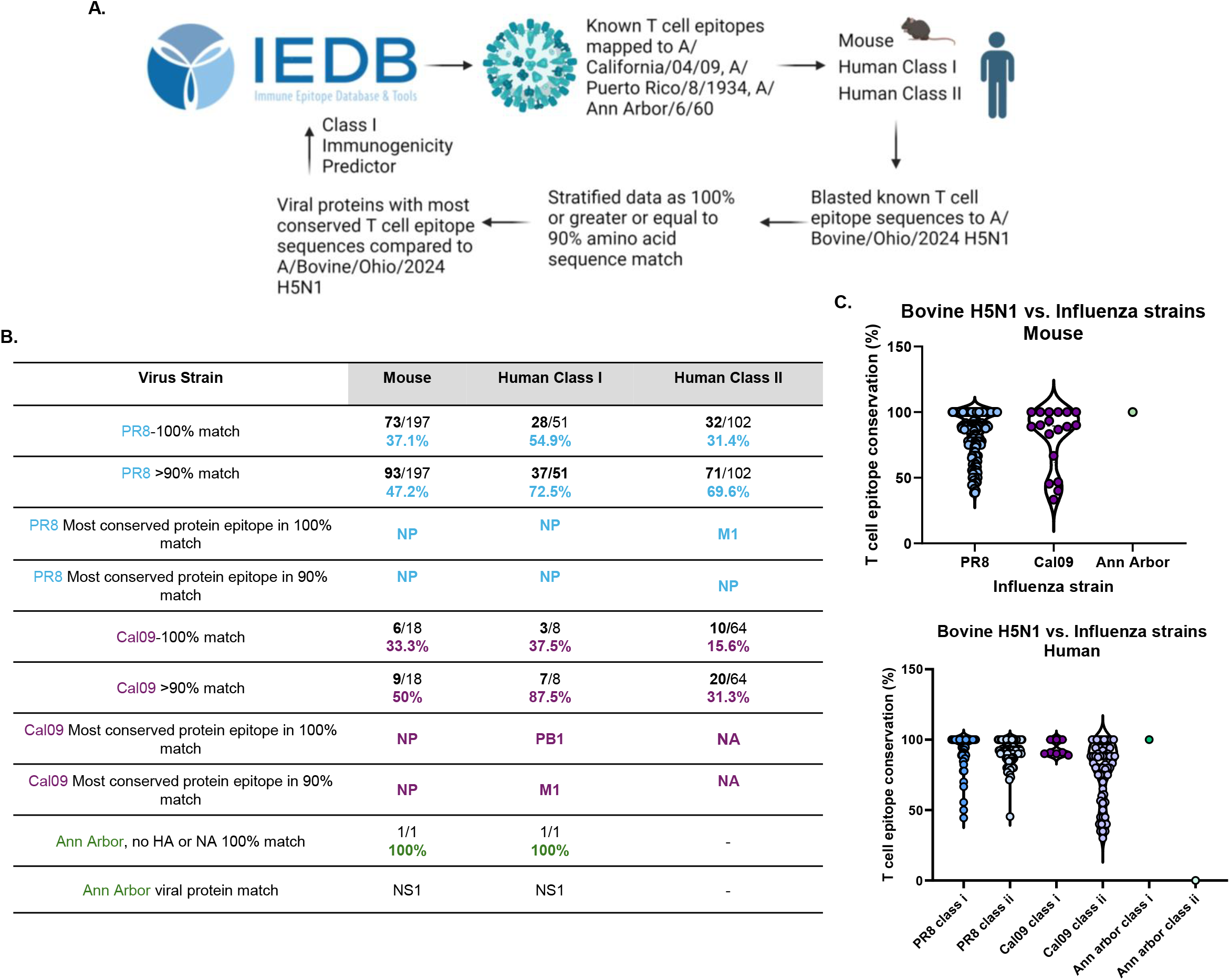
Previously mapped T cell epitopes mapped to the amino acid sequences of A/Bovine/Ohio439/2024 show conservation of T cell epitopes from H1N1 and an H2N2 strain. **(A)** Graphical summary of workflow made in *Biorender* using Immune Epitope Database and Tools **(B)** Summary table of percent of conserved T cell epitopes from PR8, Cal09, or AnnArbor60 in the A/bovine/Ohio439/2024 HPAI H5N1 virus and **(C)** Percent conserved for each known mapped T cell epitope for mouse and human, including both class I and class II restricted T cell epitopes.

For epitopes with ≥90% conservation, we observed a T cell epitope conservancy of 43.2% between PR8 and bovine H5N1 in mice, and 72.5% and 69.6% conservancy in human class I and class II, respectively (**Figure 3B and 3C**). The most conserved T cell epitopes for PR8 virus were associated with the NP viral protein (**Figure 3B, Supplemental Table 2)**. While fewer T cell epitopes were mapped for Cal09 virus, we noted similar trends to those of PR8 virus, albeit with decreased overlap in human class II epitopes (**Supplemental Tables 6, 8)**. In contrast to PR8 epitopes conserved in bovine H5N1 viruses, the most conserved epitopes across the range of viral proteins included NP, PB1, M1, and NA (**Figure 3B**).

Several T cell epitopes, including NP366 and PB1703 (*14*) are recognized as dominant correlates of protection following PR8 influenza infection in mice (*13, 16*). Furthermore, there are several immunodominant and validated human class I T cell epitopes that are important in protection against influenza challenge (*13*). Our analysis confirmed that the amino acid sequences of these immune-dominant T cell epitopes are highly conserved in the bovine HPAI H5N1 strain from both mice and human CD8 T cell epitopes (**Table 2**). Additionally, we compared the amino acid sequences of the conserved T cell epitopes (≥90% similarity) in bovine HPAI H5N1 with modern circulating H1N1 strains and those included in seasonal or 6+2 FluMist vaccines. We found a broad conservation of these sequences across H1N1 strains, ranging from 72.02% to 100%

**Table 2.**
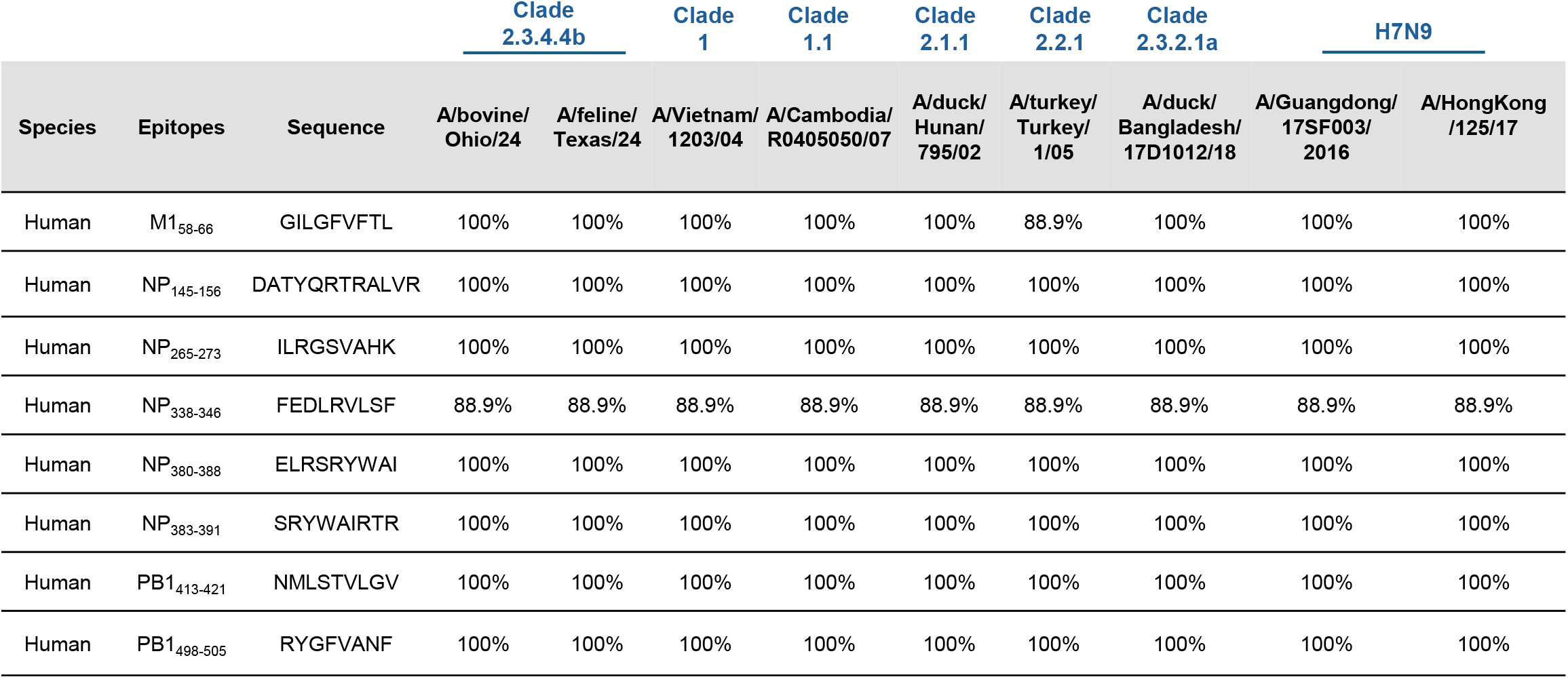
Conservation of influenza virus human CD8 T cell epitopes in the HPAI H5N1 clade 2.3.4.4b strains used in these studies (A/bovine/Ohio/24 and A/feline/NewMexico/24), other H5 clades, and H7N9 viruses.

**(Supplemental Table 1**). These findings suggest that the high conservation of known T cell epitopes from both historical and currently circulating H1N1 viruses, including immunodominant epitopes, may confer protection against HPAI H5N1 clade 2.3.4.4b viruses, in addition to the non-neutralizing antibodies we detected (**Figure 1, Supplementary Figure 2**). Collectively, these studies underscore the importance of exploring additional factors that may contribute to protection against HPAI H5N1, such as memory T cell responses, alongside traditional measures like HA inhibition or neutralizing antibodies.

## Discussion

As human infections with bovine-related HPAI H5N1 viruses continue to emerge, it is crucial to elucidate factors that mediate protection and determine the role of pre-existing influenza immunity against these viruses. Recent studies have demonstrated that prior H1N1 immunity can confer protection against bovine-derived HPAI H5N1 virus in ferrets (*9*), and mice (*11*). However, our findings indicated that neutralizing antibodies to HPAI H5N1 virus do not associate with protective immunity, highlighting the need for further investigation.

This study aimed to explore the protective effects of the pandemic H1N1 Cal09 virus and/or LAIV against HPAI bovine and cat-derived H5N1 viruses (clade 2.3.4.4b) using both murine and ferret models, with the goal of identifying potential factors that contribute to protection against lethal viral challenges. Our results show that prior infections with H1N1 viruses, including PR8 and Cal09 viruses (*11*), provided significant protection from HPAI clade 2.3.4.4b H5N1 challenges in mice. Notably, while a subset of samples did not yield positive hemagglutination inhibition or micro-neutralizing titers (data not shown), we identified cross-reactive antibodies targeting the whole H5N1 virus, predominantly directed against the HA and NA surface proteins.

Comparative amino acid sequence analysis revealed only 64.22% and 62.21% homology between the HA of HPAI H5N1 and PR8 and Cal09 viruses, respectively, with greater conservation observed in the NA protein (83.58% similarity to PR8 and 89.77% to Cal09). This suggests that the cross-protective antibodies we identified are likely directed against the NA protein but further analysis is needed. Given the high conservation of NA among H1N1 strains—including modern circulating viruses and those in seasonal influenza vaccines—it is probable that prior infection with contemporary influenza strains provides some degree of protection against newly emerged HPAI H5N1 clade 2.3.4.4b viruses.

The recent approval of the LAIV FluMist for home use in the U.S. raises questions about its effectiveness against HPAI H5N1 viruses. In our studies, we utilized three different LAIV formulations: the 2023-24 season FluMist in mice, the 2010-11 season FluMist, and a PR8 backbone LAIV containing seasonal H1N1 and H3N2 proteins under pre-clinical development at St. Jude Children’s Research Hospital. While vaccination with FluMist alone conferred mild protection against lethal HPAI H5N1 challenges in mice, the combination of LAIV with prior H1N1 infection resulted in robust protection in ferrets challenged with a cat-derived HPAI H5N1 isolate, which may possess greater crossover potential than bovine derivatives. This suggests that LAIV vaccination, especially in conjunction with pre-existing immunity, could enhance anti-H5N1 memory responses—a promising avenue for further investigation.

Memory T cell responses play a pivotal role in mediating protection to influenza viruses. We analyzed known T cell epitopes mapped for H1N1 viruses and assessed their conservation in our HPAI H5N1 strain. Our analysis revealed several conserved potential T cell epitopes, particularly in internal viral genes. Vaccination with an H5N1-specific LAIV demonstrated an increase in T cell responses to internal proteins, including matrix protein (M) and nucleoprotein (NP), thereby boosting pre-existing influenza A T cell responses from prior infections and seasonal vaccines (*12*). This may be an effective strategy, as the majority of these top conserved T cell epitopes were also found in other H5 clade viruses and H7N9 viruses (**Table 2**). These findings highlight the importance of conserved T cell memory responses in protection to HPAI H5N1 viruses.

This study has limitations. Given the urgency of the HPAI H5N1 outbreak, we utilized accessible animals, including re-purposed ferrets with existing immune histories from other ongoing lab studies, resulting in a mixed-immune history model. While we assessed pre-sera antibodies related to respective vaccine and infection histories of H1N1 strains, this data could not be incorporated into the current analysis. Another limitation is the lack of independent experiments due to time constraints and access to biosafety level 3 facilities. We recognize this limitation and ensured that we included a sufficient sample size to provide confidence in our results. In our FluMist LAIV studies, although we did not observe a significantly robust antibody response through HAI titers or HA ELISA, we did detect cross-reactive H5N1 antibodies to whole-virus and noted mild protection with decreased brain titers by 5 dpi. Future studies should utilize the most recently available LAIV to draw more definitive conclusions.

Overall, our findings identify two critical factors that may play a role in mediating protection against HPAI H5N1 in models of pre-existing immunity: non-neutralizing antibodies targeting HA and NA, and conserved memory T cell responses to highly conserved viral proteins. This research underscores the importance of understanding these immune responses to inform the development of more effective vaccines against HPAI H5N1.

## Materials and Methods

### Animal Husbandry and Ethics Statement

All procedures followed the guidelines established by the Institutional Biosafety Committee and the Animal Care and Use Committee at St. Jude Children’s Research Hospital, in accordance with the Guide for the Care and Use of Laboratory Animals as defined by the Institute of Laboratory Animal Resources and sanctioned by the U.S. National Research Council. Mice were kept under controlled conditions with 12-hour light/dark cycles at a temperature of 68°F (20°C) and 45% humidity, with continuous access to food and water. Ferrets were housed at ambient temperature (68°F (20°C) and 45% relative humidity) with 12-hour light cycles and provided species and study appropriate enrichment activities. Humane euthanasia was performed in accordance with American Veterinary Medical Association guidelines when necessary, either due to humane endpoints or for scheduled procedures. Indicators for humane endpoints included a loss of more than 30% of body weight and/or clinical scores of 3 or higher, which typically involved neurological symptoms such as circling, tremors, and reduced mobility. Ferrets were humanely euthanized via cardiac injection of Euthasol (Patterson Veterinary Supply).

### Biosafety

Experiments involving highly pathogenic avian influenza strains were conducted in a biosafety level 3 enhanced containment laboratory. Researchers wore appropriate respiratory protection (RACAL, Health and Safety Inc.) and personal protective equipment. Mice were kept in negative pressure, HEPA-filtered isolation containers. Infections or vaccinations with the pandemic California 2009 H1N1 and FluMist were conducted at biosafety level 2 prior to transferring to biosafety level 3.

### Cells, Viruses and Vaccines

Madin-Darby canine kidney (MDCK) cells (American Type Culture Collection, CCL-34) were maintained in Dulbecco’s minimum essential medium (DMEM; Corning) with 2 mM GlutaMAX (Gibco) and 10% fetal bovine serum (FBS; HyClone), at 37°C with 5% CO_2_. A/bovine/Ohio/B24OSU-439/2024 and A/California/04/2009 (H1N1) viruses were provided by Robert Webster and the Webby lab at St. Jude Children’s Research Hospital. Our lab isolated H5N1 viruses from the brain and lung of an infected cat found deceased in Curry County, New Mexico, which were used in our ferret infectivity studies (A/feline/NewMexico/C001/2024). Viruses were propagated in the allantoic cavity of 10-day-old specific-pathogen-free embryonated chicken eggs at 37°C. Allantoic fluid was clarified by centrifugation and stored at −80°C. Viral titers were assessed using 50% tissue culture infective dose (TCID_50_) assays. The following reagent was obtained through BEI Resources, NIAID, NIH: FluMist*®* Influenza Vaccine Live, Intranasal Spray, 2010-2011 Formula, NR-21987. FluMist from the 2023-2024 season was obtained from the pharmacy at St. Jude. Children’s Research Hospital.

### Mouse Lethal Dose 50 Determination

Male and female C57BL/6J mice, aged 10 to 12 weeks, were lightly anesthetized with isoflurane and intranasally inoculated with 10^1^, 10^2^, or 10^3^ TCID50 of A/Ohio/24 H5N1 in a 25 μL volume (2-4 mice per dose). Mice were monitored for 13 days for changes in body weight, clinical scores, and survival. Mice were euthanized if they lost more than 30% of body weight and/or had clinical scores of 3 or above, including neurological symptoms such as circling, tremors, and impaired motility.

### Viral titer determination

Viral titers were determined as previously described by TCID_50_ assays. Briefly, confluent MDCK cells were infected in duplicate or triplicate with 10-fold dilutions of tissue homogenates. For mice, tissues were homogenized by bead beat in 1 ml PBS, centrifuged for 5 min at 1500 x g, and supernatants were transferred to a fresh tube. For ferrets, lung tissues were finely minced with scissors before adding 1 ml of sterile PBS and bead beaten. Samples were then centrifuged for 5 min at 1500 x g to prior to measuring viral titers by TCID_50_ assay. Samples were serially diluted in minimal essential medium (MEM) plus 0.75% bovine serum albumin (BSA) and 1 μg/ml tosylsulfonyl phenylalanyl chloromethyl ketone (TPCK)-treated trypsin (H1N1 samples only). After 3 days of incubation at 37 °C and 5% CO_2_, 50 μl of the supernatant was combined and mixed with 50 μl of 0.5% packed turkey red blood cells diluted in PBS for 45 minutes at room temperature and scored by hemagglutination (HA) endpoint. Infectious viral titers were calculated using the Reed-Muench method (*17*).

### Mouse Viral Infectivity

Mice were intranasally inoculated with 10^2^ TCID50 of A/Cal/04/09, 10^5^ particles LAIV, or were naive followed by 5x mLD_50_ of A/bovine/Ohio/B24OSU-439/2024 three weeks later. They were monitored daily for 12-14 days for changes in body weight, clinical scores, and survival. Mice showing more than 30% weight loss and/or clinical scores above three were euthanized. Clinical signs were scored from 0 (no signs) to 5 (death). Neurological signs, including circling and tremors, were also observed and used as criteria for euthanasia. Post-euthanasia, lungs and brains were collected and stored at −80°C for future analysis.

### Ferret Viral Infectivity

Ferrets were lightly anesthetized using 4% inhaled isoflurane and subsequently inoculated with a virus diluted in phosphate-buffered saline (PBS) (Corning, 21-040-CV) that contained penicillin (100 U/ml) and streptomycin (100 μg/ml; Corning 30-002 CI). The viral doses administered was 10^4^ TCID_50_ of A/Feline/NewMexico/C001/2024 (H5N1). Weight and temperature were monitored daily. Clinical assessments involved a point system to evaluate the presence and severity of symptoms. Metrics included sneezing (none = 0, mild = 1, excessive = 2), coughing (absent = 0, present = 1), nasal discharge (absent = 0, present = 1), conjunctivitis (absent = 0, present = 1), and lethargy (active and playful = 0, active when stimulated = 1, not active when stimulated = 2). Mild sneezing was classified as one to two occurrences during the observation period, while continuous sneezing was labeled as excessive. Clinical evaluations were conducted by at least two researchers. Nasal washes were graded as follows: clear = 0, cloudy = 1, mucus present with discoloration = 2, and thickened mucus present = 3. Results are presented as the total score.

### Ferret nasal wash collection

Ferrets received intramuscular anesthesia using ketamine (30 mg/kg; Patterson Veterinary Supply), and sneezing was triggered by administering 1 ml of PBS containing penicillin (100 U/ml) and streptomycin (100 μg/ml) dropwise into their nostrils. Samples were collected in sterile specimen cups, briefly centrifuged, and stored at −80°C until analysis. Viral titers were assessed using the TCID_50_ assay as described above.

### Transmission Experiments

Individually housed ferrets were lightly anesthetized with 4% inhaled isoflurane and inoculated with the specified virus diluted in 500 μl of PBS. The ferrets receiving direct inoculation are referred to as index ferrets. After a 24-hour period, a contact ferret was introduced into the same cage as the index ferret, allowing for unrestricted interaction. Weight, body temperature, and clinical symptoms were monitored as previously described. Nasal washes were collected every 48 hours to assess viral transmission.

### Antibody quantification

Whole virus (H1N1 or H5N1) enzyme linked immunosorbent assays (ELISAs) were conducted using 384-well flat-bottom MaxiSorp plates (ThermoFisher) coated with either purified A/California/04/2009 at 5μg/ml or 10^6^ TCID_50_ of the virus stock of A/bovine/Ohio/B24OSU-439/2024 overnight at 4 °C. Plates were washed 4 times with PBS containing 0.1% Tween-20 (PBS-T) using the AquaMax 4000 plate washer system for H1N1 ELISA’s or handwashed for H5N1 ELISAs. Plates were blocked with PBS-T containing 0.5% Omniblok non-fat milk powder (AmericanBio) and 3% goat serum (Gibco) for 1 h at room temperature. The wash buffer was removed, and plates were tapped dry. Mouse sera was diluted 1:5 in PBS and ran in duplicate. Positive and negative mouse sera was used as controls for both sets of ELISAs. Samples were incubated at room temperature for 2 hours and then washed 4 times with PBS-T. Anti-mouse peroxidase-conjugated IgG secondary antibody was diluted at 1:3000 (Invitrogen 62-6520) in blocking buffer and 15 μL was added per well and incubated at room temperature for 1 hour. Plates were washed 4 times with PBS-T and developed using SIGMAFAST™ OPD (Sigma-Aldrich) for 10 min at room temperature. Plates were read at 490 nm using a BioTek Synergy2 plate reader and Gen5 (v3.09) software. For each plate, an upper 99% confidence interval (CI) of blank wells OD values was determined and used in determining the endpoint titers. Alternatively, mouse sera were treated with receptor-destroying enzyme (RDE; Seiken 370013), and HAI assays were performed as described (*18, 19*).

### Immune History

#### Mice

Adult female and male WT C57BL/6J mice were inoculated by IN route with ∼10^5^ particles of LAIV FluMist 2023-24 in 25 μL, or 10^2^ TCID_50_ of A/California/04/09 H1N1 in 25 μL and monitored daily for 14 days post-infection. At 3 weeks post-infection, mice were bled via eye bleeds prior to oral gavage to determine antibody titers or prior to challenge with lethal A/bovine/Ohio/B24OSU-439/2024. *Ferrets*. Neutered, de-scented male ferrets were obtained from Triple F Farms (Elmyra, NY) at the age of 6 weeks. The ferrets immune history and timeline is shown below:

| Ferret # | Sex | Vaccination | Infection Strain | Time post-Vaccination | Time post-Infection | Positive microneutralization titers? |
| --- | --- | --- | --- | --- | --- | --- |
| 4476 | Male | St. Jude LAIV | A/Cal/04/09 | 196 | 167 | Yes |
| 4477 | Male | St. Jude LAIV | A/Cal/04/09 | 196 | 167 | Yes |
| 4497 | Male | FluMist 10-11 season | A/Cal/04/09 | 196 | 167 | Yes |
| 4498 | Male | FluMist 10-11 season | A/Cal/04/09 | 196 | 167 | Yes |
| 5001 | Male | FluMist 10-11season | A/Cal/04/09 | 196 | 167 | Yes |
| 4499 | Male | St. Jude LAIV | x | 196 | 167 | Yes |
| 4500 | Male | St. Jude LAIV | x | 196 | 167 | Yes |
| 5002 | Male | St. Jude LAIV | x | 196 | 167 | Yes |

### Phylogenetic Tree Construction

H5N1 sequences were obtained from the EpiFlu database of the Global Initiative on Sharing All Influenza Data (GISAID) (https://doi.org/10.2807/1560-7917.ES.2017.22.13.30494). We downloaded all complete sequences from human, mammalian, and avian hosts in clade 2.3.4.4b from North and South America, yielding a total of 6,480 strains as of August 23, 2024. To select optimal representative sequences for phylogenetic analysis, an initial phylogenetic tree was constructed using segment HA from all downloaded strains. The large phylogeny was then down-sampled using PARNAS (*20*). The optimal representative sequences, along with our lab strains A/bovine/Ohio/B24OSU-439/2024, and A/feline/NewMexico/0002/2024 were aligned using MUSCLE (*21*) and the sequence ends were trimmed to equal lengths in AliView manually (*22*). Maximum likelihood trees were then constructed using IQ-TREE2 with the GTR+F+R5 model and 1,000 bootstrap replicates (*23*). The resulting tree files were visualized using Figtree (http://tree.bio.ed.ac.uk/software/figtree/) (24).

### Pairwise Alignment, Identity, and Mutation Detection

Pairwise amino acid sequence alignments were conducted in R with the pairwiseAlignment function from the pwalign package. Sequence identity was then determined using the pid function available at Bioconductor (https://bioconductor.org/packages/pwalign).

### T cell epitope mapping and analysis

Influenza amino acid sequences were obtained through GISIAD. Known T cell epitopes were obtained through the Immune Epitope Database and Tools (www.iedb.org)(*25*) by searching for selected influenza A virus strains with restrictions on mouse, human class I, or human class II but no disease restriction. These epitopes, viral protein information and amino acid sequence range were then used in creating tables using Microsoft Excel and the sequences were converted into FASTA files. The T cell epitopes were then compared against the known amino acid sequences for A/bovine/Ohio/B24OSU-439/2024 using the Epitope Conservancy Analysis from the Immune Epitope Database (*26*), and the percent similarity for each sequence was reported.

## Statistical Analysis

Animals were randomly selected for control and experimental groups. Some experiments had a small sample size due to mouse availability and limited access to BSL3 facilities. Graphs and statistical analyses were performed using GraphPad Prism version 10.0. Sample sizes and statistical methods, including One-way ANOVA with Tukey’s Multiple Comparisons test, Two-way ANOVA (mixed model) with multiple comparisons, a Two-way ANOVA with Kruskal-Wallis test due to repeated values, and log-rank Mantel-Cox test for survival curves, are detailed in the figure legends.

## Acknowledgements

This project has been funded in whole or in part with Federal funds from the National Institute of Allergy and Infectious Diseases, National Institutes of Health, Department of Health and Human Services, under Contract No. 75N93021C00016 (S.S.-C.), the American Lebanese Syrian Associated Charities (ALSAC) (S.S.-C.), by the National Institute of Health, Institutional Postdoctoral Training Grants (T32) Infectious Disease Therapeutics T32AI106700-08 (P.H.B.), and the National Institute of Health, Ruth L. Kirschstein Postdoctoral Individual National Research Service Award F32AI183804 (P.H.B).

The authors would like to extend their gratitude toward Richard Webby and the Webby lab at St. Jude Children’s Research Hospital for providing the original bovine isolate of HPAI H5N1 (A/bovine/Ohio.B24OSU-439/2024). We would also like to extend our gratitude to the St. Jude Children’s Research Hospital Animal Facility staff and those who operate the Biosafety Level 3 facility. Illustrations in Figures 1-3 were created with BioRender.com.

## Data availability

All data are available in the main text or the supplementary materials.

## Author contributions

Conceptualization: S.S.-C., P.H.B., B.S., E.K.R., Methodology: S.S.-C., P.H.B., E.K.R., B.L., B.S., S.T., P.G.T. Investigation: S.S.-C., P.H.B., B.S., E.K.R., V.A.M., S.T., B.L., L.L., D.R.M., T.R., K.D., V.B., P.G.T. Visualization: S.S.-C., P.H.B., S.T. Funding acquisition: S.S.-C., P.H.B. Project administration: S.S.-C., P.H.B., B.S. Supervision: S.S.-C., P.H.B., B.S. Writing – original draft: S.S.-C., P.H.B., Writing – review & editing: S.S.-C., P.H.B., V.A.M., S.T., T.R., K.D.

