## Supplemental Material for "Immune History Modifies Disease Severity to HPAI H5N1 Clade 2.3.4.4b Viral Challenge"

**Supplemental Table 1.** The mapped T cell epitopes that were conserved for A/bovine/Ohio/2024 HPAI H5N1 virus that are a greater than 90% match to modern day circulating or vaccine H1N1 strains including A/Victoria/4897/2022, A/Wisconsin/67/2022 and A/Norway/31694/2022.

| Virus Strain | Mouse | Human Class I | Human Class II |
|---|---|---|---|
| PR8 >90% match | **Victoria 2022:** 67/93, **72.02%**<br>**Wisconsin 2022:** 67/93, **72.02%**<br>**Norway 2022:** 67/93, **72.02%** | **Victoria 2022:** 29/37, **78.4%**<br>**Wisconsin 2022:** 29/37, **78.4%**<br>**Norway 2022:** 29/37, **78.4%** | **Victoria 2022:** 52/71, **73.2%**<br>**Wisconsin 2022:** 52/71, **73.2%**<br>**Norway 2022:** 53/71, **74.6%** |
| Cal09 >90% match | **Victoria 2022:** 8/9, **88.8%**<br>**Wisconsin 2022:** 8/9, **88.8%**<br>**Norway 2022:** 8/9, **88.8%** | **Victoria 2022:** 7/7, **100%**<br>**Wisconsin 2022:** 7/7, **100%**<br>**Norway 2022:** 7/7, **100%** | **Victoria 2022:** 18/20, **90%**<br>**Wisconsin 2022:** 18/20, **90%**<br>**Norway 2022:** 18/20, **90%** |
| Ann Arbor, no HA or NA 100% match | **Victoria 2022:** 0/1, **0%**<br>**Wisconsin 2022:** 0/1, **0%**<br>**Norway 2022:** 0/1, **0%** | **Victoria 2022:** 0/1, **0%**<br>**Wisconsin 2022:** 0/1, **0%**<br>**Norway 2022:** 0/1, **0%** | - |

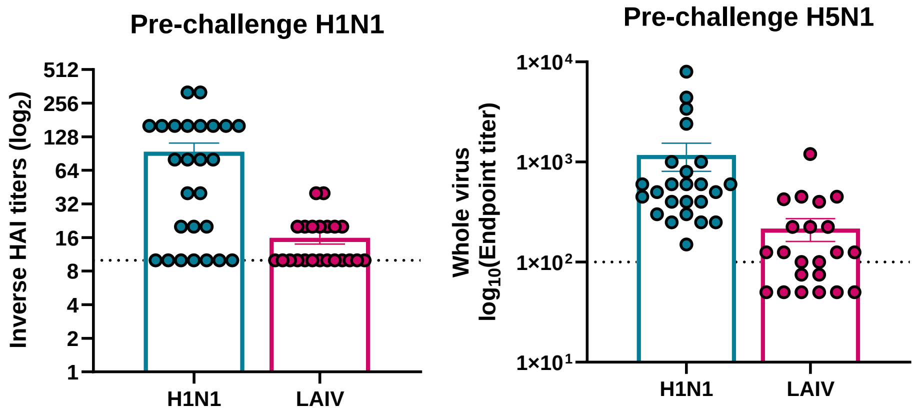


**Supplemental Figure 1.** Pre- HPAI H5N1 viral challenge sera responses to H1N1 (Cal09), LAIV (A/Victoria/2019) and HPAI bovine A/Bovine/Ohio/2024 by HAI or whole virus ELISA.

**Supplemental Table 2. Mapped PR8 T cell epitopes in mice compared to amino acid sequences from A/Bovine/Ohio/2024 H5N1 influenza virus.** Immune epitope data was obtained by the Immune Epitope Database and Tools (IEDB) program and viral sequences were obtained from GISIAD.

| Epitope name | Epitope sequence | Epitope length | Maximum identity | Starting Position | Ending Position |
|---|---|---|---|---|---|
| Non-structural protein 1 | AIMDKNIIL | 9 | **100.00%** | 122 | 130 |
| Polymerase basic protein 2 | ALLKHRFEI | 9 | **100.00%** | 70 | 78 |
| RNA-directed RNA polymerase catalytic subunit | ARLGKGYMF | 9 | **100.00%** | 349 | 357 |
| **Nucleoprotein** | **ASNENMETM** | **9** | **100.00%** | **366** | **374** |
| Nucleoprotein | AYERMCNIL | 9 | **100.00%** | 218 | 226 |
| Nucleoprotein | AYERMCNILKGK | 12 | **100.00%** | 218 | 229 |
| Neuraminidase | CVNGSCFTV | 9 | **100.00%** | 213 | 221 |
| Non-structural protein 1 | EEGAIVGEI | 9 | **100.00%** | 152 | 160 |
| Polymerase basic protein 2 | EGIPLYDA | 8 | **100.00%** | 300 | 307 |
| Polymerase basic protein 2 | ERELVRKTR | 9 | **100.00%** | 208 | 216 |
| Neuraminidase | FCGVNSDTV | 9 | **100.00%** | 430 | 438 |
| RNA-directed RNA polymerase catalytic subunit | FLEESHPGI | 9 | **100.00%** | 94 | 102 |
| polymerase PA | FLLMDALKL | 9 | **100.00%** | 282 | 290 |
| Polymerase acidic protein | FMYSDFHFI | 9 | **100.00%** | 46 | 54 |
| Nucleoprotein | FYIQMCTEL | 9 | **100.00%** | 39 | 47 |
| Nucleoprotein | GERQNATEI | 9 | **100.00%** | 17 | 25 |
| **Matrix protein 1** | **GILGFVFTL** | **9** | **100.00%** | **58** | **66** |
| Matrix protein 1 | GILGFVFTLT | 10 | **100.00%** | 58 | 67 |
| Polymerase acidic protein | GIPLYDAI | 8 | **100.00%** | 301 | 308 |
| RNA-directed RNA polymerase subunit P1 (Polymerase basic protein 1) (PB1) | GMFNMLSTV | 9 | **100.00%** | 410 | 418 |
| Nucleoprotein | IAYERMCNI | 9 | **100.00%** | 217 | 225 |
| polymerase PB2 | IGGIRMVDI | 9 | **100.00%** | 289 | 297 |
| Nucleoprotein | IGRFYIQM | 8 | **100.00%** | 36 | 43 |
| Matrix protein 1 | ILGFVFTLTV | 10 | **100.00%** | 59 | 68 |
| Matrix protein 1 | KGILGFVFTLTV | 12 | **100.00%** | 57 | 68 |
| Matrix protein 1 | KTRPILSPLTK | 11 | **100.00%** | 47 | 57 |

| | | | | | |
|---|---|---|---|---|---|
| Non-structural protein 1 | LGLDIETATRAGKQIVERI | 19 | **100.00%** | 50 | 68 |
| Polymerase basic protein 2 | LMDALKLSI | 9 | **100.00%** | 284 | 292 |
| Matrix protein 1 | MGLIYNRM | 8 | **100.00%** | 128 | 135 |
| Nuclear export protein | MITQFESL | 8 | **100.00%** | 31 | 38 |
| RNA-directed RNA polymerase catalytic subunit | MLSTVLGVSI | 10 | **100.00%** | 414 | 423 |
| Matrix protein 1 | MVLASTTAK | 9 | **100.00%** | 179 | 187 |
| RNA-directed RNA polymerase catalytic subunit | NLYNIRNLHI | 10 | **100.00%** | 597 | 606 |
| RNA-directed RNA polymerase catalytic subunit | NMLSTVLGV | 9 | **100.00%** | 413 | 421 |
| RNA-directed RNA polymerase subunit P1 (Polymerase basic protein 1) (PB1) | QPEWFRNVL | 9 | **100.00%** | 329 | 337 |
| Nucleoprotein | RFYIQMCTEL | 10 | **100.00%** | 38 | 47 |
| RNA-directed RNA polymerase catalytic subunit | RFYRTCKL | 8 | **100.00%** | 465 | 472 |
| RNA-directed RNA polymerase catalytic subunit | RLIDFLKDV | 9 | **100.00%** | 162 | 170 |
| Matrix protein 1 | RMVLASTTAK | 10 | **100.00%** | 178 | 187 |
| nucleocapsid protein | RSALILRGSVAHKSC | 15 | **100.00%** | 261 | 275 |
| RNA-directed RNA polymerase catalytic subunit | RSYLIRAL | 8 | **100.00%** | 215 | 222 |
| Nuclear export protein | RTFSFQLI | 8 | **100.00%** | 114 | 121 |
| Hemagglutinin precursor | RTLDFHDSNVK | 11 | **100.00%** | 450 | 460 |
| Neuraminidase | SGPDNGAVAV | 10 | **100.00%** | 181 | 190 |
| Neuraminidase | SGSFVQHPELTGL | 13 | **100.00%** | 388 | 400 |
| Polymerase basic protein 2 | SLENFRAYV | 9 | **100.00%** | 225 | 233 |
| Matrix protein 1 | SLLTEVETYV | 10 | **100.00%** | 2 | 11 |
| Polymerase acidic protein | SLYASPQL | 8 | **100.00%** | 648 | 655 |
| Polymerase basic protein 2 | SMIEAESSV | 9 | **100.00%** | 594 | 602 |
| **Nucleoprotein** | **SRYWAIRTR** | **9** | **100.00%** | **383** | **391** |
| Neuraminidase | SSISFCGV | 8 | **100.00%** | 426 | 433 |
| **polymerase PA** | **SSLENFRAYV** | **10** | **100.00%** | **224** | **233** |

| | | | | | |
|---|---|---|---|---|---|
| **RNA-directed RNA polymerase catalytic subunit** | **SSYRRPVGI** | **9** | **100.00%** | **703** | **711** |
| RNA-directed RNA polymerase catalytic subunit | TALANTIEV | 9 | **100.00%** | 141 | 149 |
| **Nucleoprotein** | **TYQRTRALV** | **9** | **100.00%** | **147** | **155** |
| Nucleoprotein | TYQRTRALVRTG | 12 | **100.00%** | 147 | 158 |
| nucleoprotein | TYQRTRALVRTGMDP | 15 | **100.00%** | 147 | 161 |
| Polymerase basic protein 2 | VFPNEVGARILTSE | 14 | **100.00%** | 167 | 180 |
| Polymerase basic protein 2 | VLRGFLIL | 8 | **100.00%** | 690 | 697 |
| Polymerase basic protein 2 | VYIEVLHL | 8 | **100.00%** | 227 | 234 |
| Polymerase basic protein 2 | VYINTALL | 8 | **100.00%** | 463 | 470 |
| Polymerase basic protein 2 | WMMAMKYPI | 9 | **100.00%** | 49 | 57 |
| Neuraminidase | GDVFVIREPFISCSH | 15 | **100.00%** | 97 | 111 |
| Non-structural protein 1 | IILKANFSV | 9 | **100.00%** | 128 | 136 |
| nucleoprotein | FWRGENGRRTRIAYERMCNILKG | 23 | **100.00%** | 206 | 228 |
| nucleoprotein | RLIQNSITIERMVLSAFDERRN | 22 | **100.00%** | 55 | 76 |
| neuraminidase | SISFCGV | 7 | **100.00%** | 427 | 433 |
| polymerase PA | FLLMDALKLSI | 11 | **100.00%** | 282 | 292 |
| RNA-directed RNA polymerase subunit P1 (Polymerase basic protein 1) (PB1) | GMFNMLSTVL | 10 | **100.00%** | 410 | 419 |
| RNA-directed RNA polymerase subunit P1 (Polymerase basic protein 1) (PB1) | MMGMFNMLSTV | 11 | **100.00%** | 408 | 418 |
| Neuraminidase | SGPDNGAVAVL | 11 | **100.00%** | 181 | 191 |
| Nucleoprotein | LILRGSVAHKSCL | 13 | **100.00%** | 264 | 276 |
| RNA-directed RNA polymerase subunit P1 (Polymerase basic protein 1) (PB1) | GMFNMLSTVLGVS | 13 | **100.00%** | 410 | 422 |
| Nucleoprotein | QNSQVYSLIRPNENPAHK | 18 | 94.44% | 308 | 325 |
| Nucleoprotein | IASNENMETMESSTLE | 16 | 93.75% | 365 | 380 |
| Polymerase basic protein 2 | CKISPLMVAYMLERE | 15 | 93.33% | 196 | 210 |
| Nucleoprotein | QVYSLIRPNENPAHK | 15 | 93.33% | 311 | 325 |
| Neuraminidase | TVDWSWPDGAELPFT | 15 | 93.33% | 437 | 451 |

| | | | | | |
|---|---|---|---|---|---|
| Polymerase basic protein 2 | CKISPLMVAYML | 12 | 91.67% | 196 | 207 |
| Neuraminidase | WVELIRGRPKEK | 12 | 91.67% | 408 | 419 |
| Neuraminidase | LKYNGIITETI | 11 | 90.91% | 191 | 201 |
| Hemagglutinin precursor | SVRNGTYDYPK | 11 | 90.91% | 495 | 505 |
| Nucleoprotein | TMVMELVRMIK | 11 | 90.91% | 188 | 198 |
| polymerase PB2 | CKISPLMVAYM | 11 | 90.91% | 196 | 206 |
| Nucleoprotein | ATEIRASVGK | 10 | 90.00% | 22 | 31 |
| Hemagglutinin | KEIGNGCFEF | 10 | 90.00% | 474 | 483 |
| Polymerase basic protein 2 | KISPLMVAYM | 10 | 90.00% | 197 | 206 |
| Polymerase basic protein 2 | MLLRSAIGQV | 10 | 90.00% | 548 | 557 |
| Nucleoprotein | MVMELVRMIK | 10 | 90.00% | 189 | 198 |
| Matrix protein 1 | RMGVQMQRFK | 10 | 90.00% | 243 | 252 |
| Neuraminidase | SLCPIRGWAI | 10 | 90.00% | 75 | 84 |
| Polymerase acidic R7K | SSLENFKAYV | 10 | 90.00% | 51 | 60 |
| Polymerase acidic R7T | SSLENFTAYV | 10 | 90.00% | 21 | 30 |
| Non-structural protein 1 | ALKMTMASV | 9 | 88.89% | 76 | 84 |
| Nucleoprotein S2P | APNENMETM | 9 | 88.89% | 27 | 35 |
| Nucleoprotein N3D | ASDENMETM | 9 | 88.89% | 27 | 35 |
| nucleoprotein | ASNENTETM | 9 | 88.89% | 366 | 374 |
| hemagglutinin | IYSTVASSL | 9 | 88.89% | 532 | 540 |
| Matrix protein 2 | LLTEVETPI | 9 | 88.89% | 3 | 11 |
| Hemagglutinin precursor | PVTIGECPK | 9 | 88.89% | 314 | 322 |
| Nucleoprotein | RLIQNSLTI | 9 | 88.89% | 55 | 63 |
| Matrix protein 1 | SIIPSGPLK | 9 | 88.89% | 13 | 21 |
| nucleoprotein | SPIVPSFDM | 9 | 88.89% | 473 | 481 |
| Nucleoprotein N5D | ASNEDMETM | 9 | 88.89% | 27 | 35 |
| Nucleoprotein N5H | ASNEHMETM | 9 | 88.89% | 27 | 35 |
| Nucleoprotein M9L | ASNENMETL | 9 | 88.89% | 27 | 35 |
| RNA-directed RNA polymerase subunit P1 (Polymerase basic protein 1) (PB1) | FLAMITYMT | 9 | 88.89% | 318 | 326 |
| Nuclear export protein | FMQALHLLL | 9 | 88.89% | 99 | 107 |
| nucleoprotein | LLQNSQVYS | 9 | 88.89% | 306 | 314 |
| neuraminidase NA | QIGNIISIWISHSIQTGS | 18 | 88.89% | 25 | 42 |
| RNA-directed RNA polymerase catalytic subunit | AKNMEYDA | 8 | 87.50% | 652 | 659 |

| | | | | | |
|---|---|---|---|---|---|
| Neuraminidase | CPIRGWAI | 8 | 87.50% | 77 | 84 |
| Non-structural protein 1 | FSVIFDRL | 8 | 87.50% | 134 | 141 |
| Polymerase basic protein 2 | GYEEFTMV | 8 | 87.50% | 359 | 366 |
| Neuraminidase | IRGWAIYS | 8 | 87.50% | 79 | 86 |
| RNA-directed RNA polymerase catalytic subunit | KNMEYDAV | 8 | 87.50% | 653 | 660 |
| Non-structural protein 1 | NFSVIFDR | 8 | 87.50% | 133 | 140 |
| Polymerase acidic protein | NGYIEGKL | 8 | 87.50% | 239 | 246 |
| Polymerase basic protein 2 | PNGYIEGK | 8 | 87.50% | 238 | 245 |
| Nucleoprotein | SDYEGRLI | 8 | 87.50% | 50 | 57 |
| Neuraminidase | IRGWAIYSKDNSIRI | 15 | 86.67% | 79 | 93 |
| Neuraminidase | SVAWSASACHDGMGW | 15 | 86.67% | 161 | 175 |
| Neuraminidase | ILTGNSSLCPIRGWAIYSKDN | 21 | 85.71% | 69 | 89 |
| Hemagglutinin precursor | ILAIYSTVASSL | 12 | 83.33% | 530 | 541 |
| Neuraminidase | GLASYKIFKIEKGKVTKSIELN A | 23 | 82.61% | 234 | 256 |
| Matrix protein 1 | GTHPSSSAGLK | 11 | 81.82% | 220 | 230 |
| hemagglutinin | IYSTVASSLVL | 11 | 81.82% | 532 | 542 |
| Hemagglutinin precursor | IYQILAIYSTVASSL | 15 | 80.00% | 527 | 541 |
| Hemagglutinin precursor | NVKNLYEKVK | 10 | 80.00% | 458 | 467 |
| Neuraminidase | TETIKSWRKKILRTQ | 15 | 80.00% | 198 | 212 |
| Hemagglutinin | INSSLPYQNIHPVTIGECPK | 20 | 80.00% | 302 | 321 |
| Non-structural protein 1 | LIPKQKVAGPLCIRM | 15 | 80.00% | 105 | 119 |
| Nucleoprotein | GTKVVPRGK | 9 | 77.78% | 349 | 357 |
| Hemagglutinin precursor | IEGGWTGMI | 9 | 77.78% | 354 | 362 |
| Neuraminidase | ITYKNSTWV | 9 | 77.78% | 54 | 62 |
| Matrix protein 2 | KSMREEYRK | 9 | 77.78% | 70 | 78 |
| Matrix protein 1 | RMGAVTTEV | 9 | 77.78% | 134 | 142 |
| Polymerase basic protein 2 | RMQFSSFTV | 9 | 77.78% | 630 | 638 |
| Nucleoprotein | YRRVNGKWM | 9 | 77.78% | 97 | 105 |
| neuraminidase | YRYGNGVWI | 9 | 77.78% | 336 | 344 |
| Nucleoprotein E4D/N5L | ASNDLMETM | 9 | 77.78% | 2 | 10 |
| Nucleoprotein N5H/E7A | ASNEHMATM | 9 | 77.78% | 23 | 31 |
| Nucleoprotein N5H/E7G | ASNEHMGTM | 9 | 77.78% | 27 | 35 |

| | | | | | |
|---|---|---|---|---|---|
| Nucleoprotein A1T/N5H | TSNEHMETM | 9 | 77.78% | 17 | 25 |
| hemagglutinin HA [Influenza A virus (A/Puerto Rico/8/34/Mount Sinai(H1N1))] | TICIGYHANNSTDTVDTV | 18 | 77.78% | 19 | 36 |
| Hemagglutinin precursor | IYQILAIYSTVASSLVL | 17 | 76.47% | 527 | 543 |
| Hemagglutinin precursor | FEANGNLI | 8 | 75.00% | 259 | 266 |
| Hemagglutinin precursor | MNIQFTAV | 8 | 75.00% | 403 | 410 |
| Matrix protein 2 | SLLTEVETPIRNEWGCRCNDSSDP | 24 | 75.00% | 2 | 25 |
| Hemagglutinin precursor | SSLPFQNI | 8 | 75.00% | 305 | 312 |
| matrix protein 2 | SLLTEVETPIRNEWGCRCNDSSD | 23 | 73.91% | 2 | 24 |
| Matrix protein 2 | SLLTEVETPIRNEWG | 15 | 73.33% | 2 | 16 |
| Nucleoprotein | RVLSFIKGTK | 10 | 70.00% | 342 | 351 |
| Matrix protein 2 | SLLTEVETPIRNEWGCRCNGSSD | 23 | 69.57% | 2 | 24 |
| HA1 | CPKYVRSAKLRM | 12 | 66.67% | 303 | 314 |
| Neuraminidase | GLISLILQI | 9 | 66.67% | 18 | 26 |
| Nucleoprotein | LPFDRTTVM | 9 | 66.67% | 418 | 426 |
| Protein PB1-F2 | LSLRNPILV | 9 | 66.67% | 62 | 70 |
| Hemagglutinin precursor | LYEKVKSQL | 9 | 66.67% | 462 | 470 |
| Hemagglutinin | TGLRNIPSI | 9 | 66.67% | 332 | 340 |
| Hemagglutinin precursor | VTGLRNIPS | 9 | 66.67% | 332 | 340 |
| Protein PB1-F2 | VYWKQWLSL | 9 | 66.67% | 56 | 64 |
| hemagglutinin HA [Influenza A virus (A/Puerto Rico/8/34/Mount Sinai(H1N1))] | AAADADTICIGYHANNST | 18 | 66.67% | 13 | 30 |
| Hemagglutinin precursor | IYQILAIYSTVASSLVLLVS | 20 | 65.00% | 527 | 546 |
| hypothetical 18K protein | GGLPFSLL | 8 | 62.50% | 44 | 51 |
| neuraminidase NA | SIQTGSQNHTGICNQNII | 18 | 61.11% | 37 | 54 |
| Hemagglutinin precursor | RMNYYWTLLK | 10 | 60.00% | 243 | 252 |
| Hemagglutinin | SQLKNNAKEI | 10 | 60.00% | 467 | 476 |
| Neuraminidase | VGLISLILQI | 10 | 60.00% | 17 | 26 |
| Hemagglutinin precursor | VTGLRNIPSI | 10 | 60.00% | 332 | 341 |
| Hemagglutinin precursor | ASMHECNTKCQT | 12 | 58.33% | 287 | 298 |

| | | | | | |
|---|---|---|---|---|---|
| Hemagglutinin HA | VLWGIHHPPNSK | 12 | 58.33% | 191 | 202 |
| Hemagglutinin precursor | IYQILAIYSTVASSLVLLVSLGA | 23 | 56.52% | 527 | 549 |
| Hemagglutinin HA | FERFEIFPK | 9 | 55.56% | 128 | 136 |
| Neuraminidase | TGICNQNII | 9 | 55.56% | 46 | 54 |
| neuraminidase NA | QNHTGICNQNIITYKNST | 18 | 55.56% | 43 | 60 |
| Matrix protein 1 | SLQGRTLIL | 9 | 55.56% | | |
| Matrix protein 1 | SLQGRTPIL | 9 | 55.56% | | |
| Hemagglutinin precursor | GVTAACSHAGK | 11 | 54.55% | 148 | 158 |
| Hemagglutinin precursor | VTGLRNIPSIQSR | 13 | 53.85% | 332 | 344 |
| HA1 | IVETPNSENGICY | 13 | 53.85% | 79 | 91 |
| Hemagglutinin precursor | PVRSWSYIVETPNSENGICY PGDFID | 26 | 53.85% | 89 | 114 |
| Hemagglutinin HA | RSWSYIVETPNSENGIC | 17 | 52.94% | 91 | 107 |
| Hemagglutinin HA | FERFEIFPKE | 10 | 50.00% | 128 | 137 |
| Hemagglutinin precursor | MNYYWTLL | 8 | 50.00% | 244 | 251 |
| Hemagglutinin precursor | VTAACSHAGK | 10 | 50.00% | 149 | 158 |
| Hemagglutinin precursor | VTGLRNIPSIQS | 12 | 50.00% | 332 | 343 |
| Hemagglutinin | MVTGLRNIPSIQSRGLFGAIA GFIE | 25 | 48.00% | 330 | 354 |
| Hemagglutinin precursor | NAYVSVVTSNYNRRF | 15 | 46.67% | 213 | 227 |
| Hemagglutinin precursor | EIAERPKVRDQAG | 13 | 46.15% | 230 | 242 |
| Hemagglutinin HA | SFERFEIFPKE | 11 | 45.45% | 127 | 137 |
| Hemagglutinin precursor | GIAPLQLGK | 9 | 44.44% | 63 | 71 |
| Neuraminidase | VYVDGANGV | 9 | 44.44% | 323 | 331 |
| Hemagglutinin precursor | KLKNSYVNKKGK | 12 | 41.67% | 177 | 188 |
| Hemagglutinin precursor | LTEKEGSYPK | 10 | 40.00% | 168 | 177 |
| Hemagglutinin HA | HNTNGVTAACSHE | 13 | 38.46% | 143 | 155 |
| Hemagglutinin precursor | SVSSFERFEIFPK | 13 | 38.46% | 124 | 136 |

**Supplemental Table 3. Mapped A/California/04/2009 T cell epitopes in mice compared to amino acid sequences from A/Bovine/Ohio/2024 H5N1 influenza virus.** Immune epitope data was obtained by the Immune Epitope Database and Tools (IEDB) program and viral sequences were obtained from GISIAD.

| Epitope name | Epitope sequence | Epitope length | Maximum identity | Starting Position | Ending Position |
|---|---|---|---|---|---|
| nucleocapsid protein | FYIQMCTEL | 9 | **100.00%** | 39 | 47 |
| nucleocapsid protein | RLIQNSITI | 9 | **100.00%** | 55 | 63 |
| **polymerase PB1** | **SSYRRPVGI** | **9** | **100.00%** | **703** | **711** |
| nucleocapsid protein | IQMCTELKL | 9 | **100.00%** | 41 | 49 |
| nucleocapsid protein | YSLVGIDPF | 9 | **100.00%** | 296 | 304 |
| nucleocapsid protein | RLIQNSITIERMVLS | 15 | **100.00%** | 55 | 69 |
| hemagglutinin | TYNAELLVLLENERT | 15 | 93.33% | 437 | 451 |
| polymerase PA | ASMRRNYFTA | 10 | 90.00% | 439 | 448 |
| **polymerase PA** | **PSLENFRAYV** | **10** | **90.00%** | **224** | **233** |
| nucleocapsid protein | ASNENVETM | 9 | 88.89% | 366 | 374 |
| nucleocapsid protein | TAGLTHIMI | 9 | 88.89% | 130 | 138 |
| hemagglutinin | ELLVLLENERTLDYH | 15 | 86.67% | 441 | 455 |
| matrix protein 2 | MSLLTEVETPTRSEWECRCSDSSD | 24 | 83.33% | 1 | 24 |
| hemagglutinin | TSLPFQNIHPITIGK | 15 | 66.67% | 305 | 319 |
| hemagglutinin | RYAFAMERNAGSGII | 15 | 46.67% | 269 | 283 |
| hemagglutinin | SFERFEIFPKT | 11 | 45.45% | 127 | 137 |
| hemagglutinin | SRYSKKFKPEIAIRP | 15 | 40.00% | 221 | 235 |
| hemagglutinin | KKFKPEIAIRPKVRD | 15 | 33.33% | 225 | 239 |

**Supplemental Table 4. Mapped A/Ann Arbor/60 T cell epitopes in mice compared to amino acid sequences from A/Bovine/Ohio/2024 H5N1 influenza virus.** Immune epitope data was obtained by the Immune Epitope Database and Tools (IEDB) program and viral sequences were obtained from GISIAD.

| Epitope name | Epitope sequence | Epitope length | Maximum identity | Starting Position | Ending Position |
|---|---|---|---|---|---|
| Non-structural protein 1 | AIMDKNIIL | 9 | **100.00%** | 122 | 130 |

**Supplemental Table 5. Mapped PR8 T cell epitopes for human class I restricted compared to amino acid sequences from A/Bovine/Ohio/2024 H5N1 influenza virus.** Immune epitope data was obtained by the Immune Epitope Database and Tools (IEDB) program and viral sequences were obtained from GISIAD.

| | | | | | |
|---|---|---|---|---|---|
| Non-structural protein 1 | AIMDKNIIL | 9 | **100.00%** | 122 | 130 |
| neuramindase | CVNGSCFTV | 9 | **100.00%** | 213 | 221 |
| **Nucleoprotein** | **ELRSRYWAI** | **9** | **100.00%** | **380** | **388** |
| Polymerase basic protein 2 | FMYSDFHFI | 9 | **100.00%** | 46 | 54 |
| Matrix protein 1 | GILGFVFTL | 9 | **100.00%** | 58 | 66 |
| **Matrix protein 1** | **GILGFVFTLT** | **10** | **100.00%** | **58** | **67** |
| Matrix protein 1 | ILGFVFTLTV | 10 | **100.00%** | 59 | 68 |
| **Nucleoprotein** | **ILRGSVAHK** | **9** | **100.00%** | **265** | **273** |
| **RNA-directed RNA polymerase catalytic subunit** | **NMLSTVLGV** | **9** | **100.00%** | **413** | **421** |
| Matrix protein 1 | RMVLASTTAK | 10 | **100.00%** | 178 | 187 |
| Nucleoprotein | RRSGAAGAAVK | 11 | **100.00%** | 174 | 184 |
| Hemagglutinin precursor | RTLDFHDSNVK | 11 | **100.00%** | 450 | 460 |
| Matrix protein 1 | SGPLKAEIAQRLEDV | 15 | **100.00%** | 17 | 31 |
| Polymerase basic protein 2 | SLENFRAYV | 9 | **100.00%** | 225 | 233 |
| **Nucleoprotein** | **SRYWAIRTR** | **9** | **100.00%** | **383** | **391** |
| Matrix protein 1 | TKGILGFVFTLTV | 13 | **100.00%** | 56 | 68 |
| RNA-directed RNA polymerase subunit P1 (Polymerase basic protein 1) (PB1) | VSDGGPNLY | 9 | **100.00%** | 591 | 599 |
| RNA-directed RNA polymerase subunit P1 | FVANFSMEL | 9 | **100.00%** | 501 | 509 |

| | | | | | |
|---|---|---|---|---|---|
| (Polymerase basic protein 1) (PB1) | | | | | |
| nucleoprotein | YERMCNILKG | 10 | **100.00%** | 219 | 228 |
| Polymerase basic protein 2 | SFSFGGFTFK | 10 | **100.00%** | 322 | 331 |
| nucleoprotein | HSNLNDATYQR | 11 | **100.00%** | 140 | 150 |
| nucleoprotein | IAYERMCNILKGKFQTAA | 18 | **100.00%** | 217 | 234 |
| nucleoprotein | LELRSRYWAIRTRSGGNT | 18 | **100.00%** | 379 | 396 |
| nucleoprotein | WHSNLNDATYQRTRALVR | 18 | **100.00%** | 139 | 156 |
| Polymerase basic protein 2 | SRTREILTK | 9 | **100.00%** | 14 | 22 |
| polymerase PB1 | RRAIATPGM | 9 | **100.00%** | 238 | 246 |
| **Nucleoprotein** | **DATYQRTRALVR** | **12** | **100.00%** | **145** | **156** |
| Matrix protein 1 | TRPILSPLTKGILGFVFTLTVPSERGLQRRRFV | 33 | **100.00%** | 48 | 80 |
| Matrix protein 1 | GKNTDLEVLMEWLKTRPILS | 20 | 95.00% | 34 | 53 |
| nucleoprotein | GRFYIQMCTELKLSDYEG | 18 | 94.44% | 37 | 54 |
| nucleoprotein | NGRKTRIAYERMCNILKG | 18 | 94.44% | 211 | 228 |
| nucleoprotein | NQQRASAGQISIQPTFSV | 18 | 94.44% | 397 | 414 |
| nucleoprotein | TMESSTLELRSRYWAIRT | 18 | 94.44% | 373 | 390 |
| Nucleoprotein | KLSTRGVQIASNEN | 14 | 92.86% | 357 | 370 |
| Nucleoprotein | TMVMELVRMIK | 11 | 90.91% | 188 | 198 |
| Neuraminidase | SLCPIRGWAI | 10 | 90.00% | 75 | 84 |
| nucleoprotein | GQISIQPTFS | 10 | 90.00% | 404 | 413 |
| nucleoprotein | CTELKLSDY | 9 | 88.89% | 44 | 52 |
| **Nucleoprotein** | **FEDLRVLSF** | **9** | **88.89%** | **338** | **346** |
| Matrix protein 1 | SIIPSGPLK | 9 | 88.89% | 13 | 21 |
| nucleoprotein | AAFEDLRVL | 9 | 88.89% | 336 | 344 |
| Nucleoprotein | SAAFEDLRVLSFIKG | 15 | 86.67% | 335 | 349 |
| nucleocapsid protein | PKKTGGPIYRRVN | 13 | 84.62% | 89 | 101 |
| Matrix protein 1 | RGLQRRRFVQNALNGNG | 17 | 82.35% | 72 | 88 |
| Matrix protein 2 | KSMREEYRK | 9 | 77.78% | 70 | 78 |
| nucleoprotein | LPFDRTTIM | 9 | 77.78% | 418 | 426 |
| Nucleoprotein | RVLSFIKGTK | 10 | 70.00% | 342 | 351 |
| Matrix protein 2 | VETPIRNEW | 9 | 66.67% | 7 | 15 |
| Hemagglutinin precursor | GIHHPSNSK | 9 | 55.56% | 195 | 203 |
| Hemagglutinin precursor | VTAACSHAGK | 10 | 50.00% | 149 | 158 |
| Hemagglutinin precursor | GIAPLQLGK | 9 | 44.44% | 63 | 71 |

**Supplemental Table 6. Mapped PR8 T cell epitopes for human class II restricted compared to amino acid sequences from A/Bovine/Ohio/2024 H5N1 influenza virus.** Immune epitope data was obtained by the Immune Epitope Database and Tools (IEDB) program and viral sequences were obtained from GISIAD.

| Epitope name | Epitope sequence | Epitope length | Maximum identity | Starting Position | Ending Position |
|---|---|---|---|---|---|
| nucleocapsid protein | DATYQRTRALVRTGMDPRMC | 20 | **100.00%** | 145 | 164 |
| Matrix protein 1 | FVFTLTVPSER | 11 | **100.00%** | 62 | 72 |
| nucleocapsid protein | MIWHSNLNDATYQRTRALVR | 20 | **100.00%** | 137 | 156 |
| Matrix protein 1 | SGPLKAEIAQRLEDV | 15 | **100.00%** | 17 | 31 |
| matrix protein 1 | TNPLIRHENRMVLASTTAKA | 20 | **100.00%** | 169 | 188 |
| nucleoprotein | ATYQRTRALVRTG | 13 | **100.00%** | 146 | 158 |
| matrix protein 1 | GPLKAEIAQRLED | 13 | **100.00%** | 18 | 30 |
| matrix protein 1 | LGFVFTLTVPSER | 13 | **100.00%** | 60 | 72 |
| nucleoprotein | NKYLEEHPSAGKD | 13 | **100.00%** | 76 | 88 |
| matrix protein 1 | NPLIRHENRMVLAS | 14 | **100.00%** | 170 | 183 |
| matrix protein 1 | QAMRTIGTHPSSS | 13 | **100.00%** | 214 | 226 |
| matrix protein 1 | QMVQAMRTIGTHP | 13 | **100.00%** | 211 | 223 |
| matrix protein 1 | RKLKREITFHGAK | 13 | **100.00%** | 101 | 113 |
| matrix protein 1 | RMVLASTTAKAMEQ | 14 | **100.00%** | 178 | 191 |
| matrix protein 1 | DLLENLQAYQKRM | 13 | **100.00%** | 232 | 244 |
| nucleoprotein | ELILYDKEEIRRI | 13 | **100.00%** | 107 | 119 |

| | | | | | |
|---|---|---|---|---|---|
| nucleoprotein | ILYDKEEIRRIWRQ | 14 | **100.00 %** | 109 | 122 |
| matrix protein 1 | NPLIRHENRMVLA | 13 | **100.00 %** | 170 | 182 |
| matrix protein 1 | PLIRHENRMVLAST | 14 | **100.00 %** | 171 | 184 |
| matrix protein 1 | RQMVQAMRTIGTH | 13 | **100.00 %** | 210 | 222 |
| matrix protein 1 | EAMEVASQARQMVQAMRTIG | 20 | **100.00 %** | 201 | 220 |
| nucleocapsid protein | ALVRTGMDPRMCSLMQGSTL | 20 | **100.00 %** | 153 | 172 |
| nucleocapsid protein | MRELILYDKEEIRRIWRQAN | 20 | **100.00 %** | 105 | 124 |
| nucleoprotein | STLELRSRYWAIRTRSGGNT | 20 | **100.00 %** | 377 | 396 |
| matrix protein M1 | ARQMVQAMRTIGT | 13 | **100.00 %** | 209 | 221 |
| nucleoprotein NP | GKWMRELILYDKE | 13 | **100.00 %** | 102 | 114 |
| matrix protein M1 | MEWLKTRPILSPL | 13 | **100.00 %** | 43 | 55 |
| matrix protein M1 | RKLKREITFHGA | 12 | **100.00 %** | 101 | 112 |
| RNA-directed RNA polymerase subunit P1 (Polymerase basic protein 1) (PB1) | GMFNMLSTVLGVS | 13 | **100.00 %** | 410 | 422 |
| RNA-directed RNA polymerase catalytic subunit | NVLSIAPIMFSNKMARLGKG | 20 | **100.00 %** | 335 | 354 |
| RNA-directed RNA polymerase catalytic subunit | NVVRKMMTNSQDTELSFTIT | 20 | **100.00 %** | 284 | 303 |
| RNA-directed RNA polymerase subunit P1 (Polymerase basic protein 1) (PB1) | PGMMMGMFNMLSTVLGVSIL | 20 | **100.00 %** | 405 | 424 |
| nucleoprotein | CNILKGKFQTAAQKAMMDQVRESR | 24 | 95.83% | 223 | 246 |
| nucleoprotein | ASAGQISIQPTFSVQRNLPF | 20 | 95.00% | 401 | 420 |
| nucleocapsid protein | GVGTMVMELVRMIKRGINDR | 20 | 95.00% | 185 | 204 |
| matrix protein 1 | TYVLSIIPSGPLKAEIAQRL | 20 | 95.00% | 9 | 28 |
| Matrix protein 1 | VLMEWLKTRPILSPLTKGIL | 20 | 95.00% | 41 | 60 |
| matrix protein 1 | DPNNMDKAVKLYRKLKREIT | 20 | 95.00% | 89 | 108 |
| matrix protein 1 | MSLLTEVETYVLSIIPSGPL | 20 | 95.00% | 1 | 20 |
| nucleocapsid protein | KEEIRRIWRQANNGDDATAG | 20 | 95.00% | 113 | 132 |

| | | | | | |
|---|---|---|---|---|---|
| Matrix protein 1 | AGKNTDLEVLMEWLKTRPIL | 20 | 95.00% | 33 | 52 |
| nucleoprotein | IASNENMETMESSTL | 15 | 93.33% | 365 | 379 |
| nucleoprotein | GTMVMELVRMIKRGI | 15 | 93.33% | 187 | 201 |
| nucleoprotein | EGRLIQNSLTIERMV | 15 | 93.33% | 53 | 67 |
| nucleoprotein | GERQNATEIRASVGK | 15 | 93.33% | 17 | 31 |
| Nucleoprotein S2P | LGSRYWAIRTRSGGN | 15 | 93.33% | 42 | 56 |
| nucleoprotein | PACVYGPAVASGYDF | 15 | 93.33% | 277 | 291 |
| nucleoprotein | RTEIIRMMESARPED | 15 | 93.33% | 441 | 455 |
| nucleoprotein | TFLARSALILRGSVA | 15 | 93.33% | 257 | 271 |
| nucleoprotein | TSDMRTEIIRMMESA | 15 | 93.33% | 437 | 451 |
| nucleoprotein | GQISIQPTFSVQRN | 14 | 92.86% | 404 | 417 |
| nucleoprotein | RLIQNSLTIERMVL | 14 | 92.86% | 55 | 68 |
| nucleoprotein | DPFRLLQNSQVYS | 13 | 92.31% | 302 | 314 |
| nucleoprotein | EIIRMMESARPED | 13 | 92.31% | 443 | 455 |
| nucleoprotein | RMMESARPEDVSF | 13 | 92.31% | 446 | 458 |
| matrix protein 1 | VLMEWLKTRPILS | 13 | 92.31% | 41 | 53 |
| nucleoprotein NP | GQISIQPTFSVQR | 13 | 92.31% | 404 | 416 |
| matrix protein 1 | YQKRMGVQMQRFK | 13 | 92.31% | 240 | 252 |
| matrix protein M1 | DKAVKLYRKLKRE | 13 | 92.31% | 94 | 106 |
| nucleoprotein NP | EIRRIWRQANNGD | 13 | 92.31% | 115 | 127 |
| nucleoprotein | GRGVFELSDEKAA | 13 | 92.31% | 460 | 472 |
| nucleoprotein | QPTFSVQRNLPFD | 13 | 92.31% | 409 | 421 |
| nucleoprotein | RQNATEIRASVGKMIGGIGR FYIQ | 24 | 91.67% | 19 | 42 |
| Nucleoprotein | EDLTFLARSAL | 11 | 90.91% | 254 | 264 |
| matrix protein 1 | VLMEWLKTRPI | 11 | 90.91% | 41 | 51 |
| nucleoprotein | QVRESRNPGNAEFEDLTFLA | 20 | 90.00% | 241 | 260 |
| matrix protein 1 | SHRQMVTTTNPLIRHENRMV | 20 | 90.00% | 161 | 180 |
| matrix protein 1 | VKLYRKLKREITFHGAKEIS | 20 | 90.00% | 97 | 116 |
| nucleocapsid protein | LSDYEGRLIQNSLTIERMVL | 20 | 90.00% | 49 | 68 |
| Nucleoprotein | QPTFSVQRNLPFDRTTIMAA | 20 | 90.00% | 409 | 428 |
| Nucleoprotein | LPRGKLSTRGVQIASNENME | 20 | 90.00% | 353 | 372 |
| nucleoprotein | EKAASPIVPSFDMSNEGSYF FGDN | 24 | 87.50% | 469 | 492 |
| nucleocapsid protein | AAQKAMMDQVRESRD | 15 | 86.67% | 233 | 247 |
| Nucleoprotein S2P | ENMETMESSTLELGS | 15 | 86.67% | 30 | 44 |
| Nucleoprotein S2P | IAPNENMETMESSTL | 15 | 86.67% | 26 | 40 |
| Nucleoprotein N5D | IASNEDMETMESSTL | 15 | 86.67% | 26 | 40 |

| | | | | | |
|---|---|---|---|---|---|
| Nucleoprotein N5H | IASNEHMETMESSTL | 15 | 86.67% | 26 | 40 |
| Nucleoprotein | ITSNENMETMESSTL | 15 | 86.67% | | |
| Nucleoprotein N3D | KLSTRGVQIASDENM | 15 | 86.67% | 18 | 32 |
| Nucleoprotein N5D | KLSTRGVQIASNEDM | 15 | 86.67% | 18 | 32 |
| Nucleoprotein S2P | TMESSTLELGSRYWA | 15 | 86.67% | 34 | 48 |
| nucleoprotein | YRRVNGKWMRELILY | 15 | 86.67% | 97 | 111 |
| matrix protein 1 | GLQRRRFVQNALNGNGDPNN | 20 | 85.00% | 73 | 92 |
| matrix protein 1 | SSAGLKNDLLENLQAYQKRM | 20 | 85.00% | 225 | 244 |
| matrix protein M1 | REITFHGAKEISL | 13 | 84.62% | 105 | 117 |
| matrix protein 1 | RQMVTTTNPLIRH | 13 | 84.62% | 163 | 175 |
| matrix protein M1 | GLIYNRMGAVTTE | 13 | 84.62% | 129 | 141 |
| nucleoprotein NP | VFELSDEKAASPI | 13 | 84.62% | 463 | 475 |
| nucleoprotein | AFEDLRVLSFIKGTKVLPRG | 20 | 80.00% | 337 | 356 |
| Nucleoprotein N5H/E7A | GKLSNRGVQIASNEH | 15 | 80.00% | 13 | 27 |
| Nucleoprotein N5H/E7G | IASNEHMGTMESSTL | 15 | 80.00% | 26 | 40 |
| nucleoprotein | NLPFDRTTIMAAFNG | 15 | 80.00% | 417 | 431 |
| nucleocapsid protein | NLPFDRTTVMAAFTG | 15 | 80.00% | 417 | 431 |
| Nucleoprotein N5H/E7G | RGVQIASNEHMGTME | 15 | 80.00% | 22 | 36 |
| Nucleoprotein N5H/E7A | VLPRGKLSNRGVQIA | 15 | 80.00% | 9 | 23 |
| Matrix protein 1 | ITFHGAKEIS | 10 | 80.00% | 107 | 116 |
| matrix protein 1 | RRFVQNALNGNGDP | 14 | 78.57% | 77 | 90 |
| matrix protein 1 | KEISLSYSAGALA | 13 | 76.92% | 113 | 125 |
| Nucleoprotein N5H/E7A | NRGVQIASNEHMATME | 16 | 75.00% | 17 | 32 |
| nucleoprotein [Influenza A virus (A/Puerto Rico/8/1934(H1N1))] | NLPFDRTTVMAAFSG | 15 | 73.33% | 417 | 431 |
| nucleoprotein | RVLSFIKGTKVLPR | 14 | 71.43% | 342 | 355 |
| Hemagglutinin precursor | SFERFEIFPKE | 11 | 45.45% | 127 | 137 |

**Supplemental Table 7. Mapped A/California/04/09 T cell epitopes for human class I restricted compared to amino acid sequences from A/Bovine/Ohio/2024 H5N1 influenza virus.** Immune epitope data was obtained by the Immune Epitope Database and Tools (IEDB) program and viral sequences were obtained from GISIAD.

| Epitope name | Epitope sequence | Epitope length | Maximum identity | Starting Position | Ending Position |
|---|---|---|---|---|---|
| **polymerase PB1** | **RYGFVANF** | **8** | **100.00%** | **498** | **505** |
| **polymerase PB1** | **FYRYGFVANF** | **10** | **100.00%** | **496** | **505** |
| polymerase PA | LYASPQLEGF | 10 | **100.00%** | 649 | 658 |
| matrix protein 1 | LYKKLKREITF | 11 | 90.91% | 99 | 109 |
| nucleocapsid protein | PFERATVMAAF | 11 | 90.91% | 419 | 429 |
| matrix protein 1 | AYQKRMGVQM | 10 | 90.00% | 239 | 248 |
| matrix protein 1 | TFHGAKEVSL | 10 | 90.00% | 108 | 117 |
| nucleocapsid protein | LPFERATVM | 9 | 88.89% | 418 | 426 |

**Supplemental Table 8. Mapped A/California/04/09 T cell epitopes for human class II restricted compared to amino acid sequences from A/Bovine/Ohio/2024 H5N1 influenza virus.** Immune epitope data was obtained by the Immune Epitope Database and Tools (IEDB) program and viral sequences were obtained from GISIAD.

| Epitope name | Epitope sequence | Epitope length | Maximum identity | Starting Position | Ending Position |
|---|---|---|---|---|---|
| neuraminidase | CFWVELIRGRPKENTIW | 17 | **100.00%** | 421 | 437 |
| neuraminidase | DNGAVAVLKYNGIITDT | 17 | **100.00%** | 199 | 215 |
| neuraminidase | GSKGDVFVIREPFISCS | 17 | **100.00%** | 109 | 125 |
| neuraminidase | IITDTIKSWRNNILRTQ | 17 | **100.00%** | 211 | 227 |
| neuraminidase | ILRTQESECACVNGSCF | 17 | **100.00%** | 223 | 239 |
| neuraminidase | KSWRNNILRTQESECAC | 17 | **100.00%** | 217 | 233 |
| neuraminidase | LECRTFFLTQGALLNDK | 17 | **100.00%** | 127 | 143 |
| neuraminidase | NQNLEYQIGYICSGIFG | 17 | **100.00%** | 307 | 323 |
| neuraminidase | VLKYNGIITDTIKSWRN | 17 | **100.00%** | 205 | 221 |
| matrix protein 1 | GFVFTLTVPSE | 11 | **100.00%** | 61 | 71 |
| matrix protein 1 | TYVLSIIPSGPLKAEIAQRL | 20 | 95.00% | 9 | 28 |
| Nucleoprotein | NPAHKSQLVWMACHSAAFEI | 20 | 95.00% | | |
| hemagglutinin | WTYNAELLVLLENERTLD | 18 | 94.44% | 436 | 453 |
| neuraminidase | EKGKIVKSVEMNAPNYH | 17 | 94.12% | 259 | 275 |
| neuraminidase | FSFKYGNGVWIGRTKSI | 17 | 94.12% | 349 | 365 |

| | | | | | |
|---|---|---|---|---|---|
| neuraminidase | FVIREPFISCSPLECRT | 17 | 94.12% | 115 | 131 |
| neuraminidase | LDCIRPCFWVELIRGRP | 17 | 94.12% | 415 | 431 |
| matrix protein 1 | KKLKREITFHGAK | 13 | 92.31% | 101 | 113 |
| hemagglutinin | ELLVLLENERTLDYHDSNVK | 20 | 90.00% | 441 | 460 |
| hemagglutinin | LDIWTYNAELLVLLENERTL | 20 | 90.00% | 433 | 452 |
| hemagglutinin | TGMVDGWYGYHHQNEQGS | 18 | 88.89% | 359 | 376 |
| hemagglutinin | ELLVLLENERTLDYHDS | 17 | 88.24% | 441 | 457 |
| neuraminidase | GQASYKIFRIEKGKIVK | 17 | 88.24% | 249 | 265 |
| neuraminidase | GWAIYSKDNSVRIGSKG | 17 | 88.24% | 96 | 112 |
| neuraminidase | NGVWIGRTKSISSRNGF | 17 | 88.24% | 355 | 371 |
| neuraminidase | QIGNIISIWISHSIQLG | 17 | 88.24% | 25 | 41 |
| neuraminidase | SNGQASYKIFRIEKGKI | 17 | 88.24% | 247 | 263 |
| neuraminidase | SRNGFEMIWDPNGWTGT | 17 | 88.24% | 367 | 383 |
| neuraminidase | WAIYSKDNSVRIGSKGD | 17 | 88.24% | 97 | 113 |
| neuraminidase | YKIFRIEKGKIVKSVEM | 17 | 88.24% | 253 | 269 |
| neuraminidase | GFEMIWDPNGWTGTDN | 16 | 87.50% | 370 | 385 |
| hemagglutinin | ILAIYSTVASSL | 12 | 83.33% | 530 | 541 |
| hemagglutinin | NKVNSVIEKMNTQFTAVG | 18 | 83.33% | 394 | 411 |
| neuraminidase | MNPNQKIITIGSVCMTI | 17 | 82.35% | 1 | 17 |
| hemagglutinin | SNVKNLYEKVRSQLKNNAKE | 20 | 80.00% | 457 | 476 |
| hemagglutinin | AVGKEFNHLEKRIENLNKKV | 20 | 80.00% | 409 | 428 |
| hemagglutinin | HPITIGKCPKYVKSTKLRLA | 20 | 80.00% | 313 | 332 |
| hemagglutinin | KVRSQLKNNAKEIGNGCFEF | 20 | 80.00% | 465 | 484 |
| hemagglutinin | PKYVKSTKLRLATGLRNIPS | 20 | 80.00% | 321 | 340 |
| hemagglutinin | TNKVNSVIEKMNTQFTAVGK | 20 | 80.00% | 393 | 412 |
| nucleocapsid protein | RTEVIRMMESAKPEDLSFQ | 19 | 78.95% | 441 | 459 |
| hemagglutinin | NLYEKVRSQLKNNAKEIGNG | 20 | 75.00% | 461 | 480 |

| | | | | | |
|---|---|---|---|---|---|
| hemagglutinin | EKMNTQFTAVGKEFNHLEKR | 20 | 75.00% | 401 | 420 |
| hemagglutinin | KLESTRIYQILAIYSTVASS | 20 | 75.00% | 521 | 540 |
| hemagglutinin | TSLPFQNIHPITIGKCPKYV | 20 | 75.00% | 305 | 324 |
| hemagglutinin | NREEIDGVKLESTRIYQILA | 20 | 70.00% | 513 | 532 |
| hemagglutinin | VDTVLEKNVTVTHSVNLLED | 20 | 65.00% | 33 | 52 |
| hemagglutinin | VTHSVNLLEDKHNGKLCK | 18 | 61.11% | 43 | 60 |
| hemagglutinin | KLCKLRGVAPLHLGKCNIAG | 20 | 60.00% | 57 | 76 |
| hemagglutinin | IYQILAIYSTVASSLVLVVSLGA | 23 | 56.52% | 527 | 549 |
| hemagglutinin | VLEKNVTVTHSVNLLEDK | 18 | 55.56% | 36 | 53 |
| hemagglutinin | LRLATGLRNIPSIQSRGLFG | 20 | 55.00% | 329 | 348 |
| hemagglutinin | EGRMNYYWTLVEPGDKITFE | 20 | 50.00% | 241 | 260 |
| hemagglutinin | IDYEELREQLSSVSSFERFE | 20 | 45.00% | 113 | 132 |
| hemagglutinin | KKFKPEIAIRPKVRDQEGRM | 20 | 45.00% | 225 | 244 |
| hemagglutinin | TLVEPGDKITFEATGNLVVP | 20 | 45.00% | 249 | 268 |
| hemagglutinin | QLSSVSSFERFEIFPKTSSW | 20 | 40.00% | 121 | 140 |
| hemagglutinin | FVGSSRYSKKFKPEIAIRPK | 20 | 40.00% | 217 | 236 |
| hemagglutinin | FYKNLIWLVKKGNSYPKLSK | 20 | 40.00% | 161 | 180 |
| hemagglutinin | ITFEATGNLVVPRYAFAMER | 20 | 40.00% | 257 | 276 |
| hemagglutinin | ERFEIFPKTSSWPNHDSNKG | 20 | 35.00% | 129 | 148 |
| hemagglutinin | YQNADTYVFVGSSRYSKKFK | 20 | 35.00% | 209 | 228 |
| hemagglutinin | LVVPRYAFAMERNAGSGIII | 20 | 35.00% | 265 | 284 |
| hemagglutinin | MKAILVVLLYTFATANADTL | 20 | 30.00% | 1 | 20 |

**Supplemental Table 9. T cell epitopes from Cal09 human class I restricted that were a 90 or greater percent match to bovine H5N1 conserved in Victoria, Wisconsin, or Norway H1N1 2022.** Immune epitope data was obtained by the Immune Epitope Database and Tools (IEDB) program and viral sequences were obtained from GISIAD.

| Epitope name | Epitope sequence | Epitope length | Maximum identity | Starting Position | Ending Position |
|---|---|---|---|---|---|
| **A/Victoria/4897/2022** | | | | | |
| **polymerase PB1** | **RYGFVANF** | **8** | **100.00%** | **498** | **505** |
| **polymerase PB1** | **FYRYGFVANF** | **10** | **100.00%** | **496** | **505** |
| polymerase PA | LYASPQLEGF | 10 | **100.00%** | 649 | 658 |
| matrix protein 1 | LYKKLKREITF | 11 | **100.00%** | 99 | 109 |
| matrix protein 1 | AYQKRMGVQM | 10 | **100.00%** | 239 | 248 |
| matrix protein 1 | TFHGAKEVSL | 10 | **100.00%** | 108 | 117 |
| nucleocapsid protein | PFERATVMAAF | 11 | 90.91% | 419 | 429 |
| **A/Wisconsin/67/2022** | | | | | |
| **polymerase PB1** | **RYGFVANF** | **8** | **100.00%** | **498** | **505** |
| **polymerase PB1** | **FYRYGFVANF** | **10** | **100.00%** | **496** | **505** |
| polymerase PA | LYASPQLEGF | 10 | **100.00%** | 649 | 658 |
| matrix protein 1 | LYKKLKREITF | 11 | **100.00%** | 99 | 109 |
| matrix protein 1 | AYQKRMGVQM | 10 | **100.00%** | 239 | 248 |
| matrix protein 1 | TFHGAKEVSL | 10 | **100.00%** | 108 | 117 |
| nucleocapsid protein | PFERATVMAAF | 11 | 90.91% | 419 | 429 |
| **A/Norway/31694/2022** | | | | | |
| **polymerase PB1** | **RYGFVANF** | **8** | **100.00%** | **498** | **505** |
| **polymerase PB1** | **FYRYGFVANF** | **10** | **100.00%** | **496** | **505** |

| | | | | | |
|---|---|---|---|---|---|
| polymerase PA | LYASPQLEGF | 10 | **100.00%** | 649 | 658 |
| matrix protein 1 | LYKKLKREITF | 11 | **100.00%** | 99 | 109 |
| matrix protein 1 | AYQKRMGVQM | 10 | **100.00%** | 239 | 248 |
| matrix protein 1 | TFHGAKEVSL | 10 | **100.00%** | 108 | 117 |
| nucleocapsid protein | PFERATVMAAF | 11 | 90.91% | 419 | 429 |

**Supplemental Table 10. T cell epitopes from Cal09 human class II restricted that were a 90 or greater percent match to bovine H5N1 conserved in Victoria, Wisconsin, or Norway H1N1 2022.** Immune epitope data was obtained by the Immune Epitope Database and Tools (IEDB) program and viral sequences were obtained from GISIAD.

| Epitope name | Epitope sequence | Epitope length | Maximum identity | Starting Position | Ending Position |
|---|---|---|---|---|---|
| **A/Victoria/4897/2022** | | | | | |
| neuraminidase | GSKGDVFVIREPFISCS | 17 | **100.00%** | 436 | 453 |
| neuraminidase | ILRTQESECACVNGSCF | 17 | **100.00%** | 101 | 113 |
| neuraminidase | LECRTFFLTQGALLNDK | 17 | **100.00%** | 433 | 452 |
| neuraminidase | VLKYNGIITDTIKSWRN | 17 | **100.00%** | 211 | 227 |
| matrix protein 1 | GFVFTLTVPSE | 11 | **100.00%** | 415 | 431 |
| matrix protein 1 | TYVLSIIPSGPLKAEIAQRL | 20 | **100.00%** | 307 | 323 |
| hemagglutinin | WTYNAELLVLLENERTLD | 18 | **100.00%** | 259 | 275 |
| neuraminidase | FSFKYGNGVWIGRTKSI | 17 | **100.00%** | 223 | 239 |
| neuraminidase | FVIREPFISCSPLECRT | 17 | **100.00%** | 205 | 221 |
| matrix protein 1 | KKLKREITFHGAK | 13 | **100.00%** | 217 | 233 |
| hemagglutinin | ELLVLLENERTLDYHDSNVK | 20 | **100.00%** | 349 | 365 |
| Nucleoprotein | NPAHKSQLVWMACHSAAFEI | 20 | 95.00% | | |
| hemagglutinin | LDIWTYNAELLVLLENERTL | 20 | 95.00% | 421 | 437 |
| neuraminidase | CFWVELIRGRPKENTIW | 17 | 94.12% | 127 | 143 |
| neuraminidase | DNGAVAVLKYNGIITDT | 17 | 94.12% | 61 | 71 |
| neuraminidase | IITDTIKSWRNNILRTQ | 17 | 94.12% | 115 | 131 |
| neuraminidase | KSWRNNILRTQESECAC | 17 | 94.12% | 441 | 460 |
| neuraminidase | LDCIRPCFWVELIRGRP | 17 | 94.12% | 9 | 28 |

| | | | | | |
|---|---|---|---|---|---|
| neuraminidase | NQNLEYQIGYICSGIFG | 17 | 88.24% | 199 | 215 |
| neuraminidase | EKGKIVKSVEMNAPNYH | 17 | 88.24% | 109 | 125 |
| **A/Wisconsin/67/2022** | | | | | |
| neuraminidase | GSKGDVFVIREPFISCS | 17 | **100.00%** | 127 | 143 |
| neuraminidase | ILRTQESECACVNGSCF | 17 | **100.00%** | 61 | 71 |
| neuraminidase | LECRTFFLTQGALLNDK | 17 | **100.00%** | 436 | 453 |
| neuraminidase | VLKYNGIITDTIKSWRN | 17 | **100.00%** | 115 | 131 |
| matrix protein 1 | GFVFTLTVPSE | 11 | **100.00%** | 101 | 113 |
| matrix protein 1 | TYVLSIIPSGPLKAEIAQRL | 20 | **100.00%** | 441 | 460 |
| hemagglutinin | WTYNAELLVLLENERTLD | 18 | **100.00%** | 433 | 452 |
| neuraminidase | FSFKYGNGVWIGRTKSI | 17 | **100.00%** | 199 | 215 |
| neuraminidase | FVIREPFISCSPLECRT | 17 | **100.00%** | 211 | 227 |
| matrix protein 1 | KKLKREITFHGAK | 13 | **100.00%** | 415 | 431 |
| hemagglutinin | ELLVLLENERTLDYHDSNVK | 20 | **100.00%** | 307 | 323 |
| Nucleoprotein | NPAHKSQLVWMACHSAAFEI | 20 | 95.00% | | |
| hemagglutinin | LDIWTYNAELLVLLENERTL | 20 | 95.00% | 259 | 275 |
| neuraminidase | CFWVELIRGRPKENTIW | 17 | 94.12% | 109 | 125 |
| neuraminidase | DNGAVAVLKYNGIITDT | 17 | 94.12% | 223 | 239 |
| neuraminidase | IITDTIKSWRNNILRTQ | 17 | 94.12% | 205 | 221 |
| neuraminidase | KSWRNNILRTQESECAC | 17 | 94.12% | 9 | 28 |
| neuraminidase | LDCIRPCFWVELIRGRP | 17 | 94.12% | 217 | 233 |
| neuraminidase | NQNLEYQIGYICSGIFG | 17 | 88.24% | 349 | 365 |
| neuraminidase | EKGKIVKSVEMNAPNYH | 17 | 88.24% | 421 | 437 |
| **A/Norway/31694/2022** | | | | | |
| neuraminidase | GSKGDVFVIREPFISCS | 17 | 100.00% | 109 | 125 |
| neuraminidase | ILRTQESECACVNGSCF | 17 | 100.00% | 223 | 239 |
| neuraminidase | LECRTFFLTQGALLNDK | 17 | 100.00% | 127 | 143 |
| neuraminidase | VLKYNGIITDTIKSWRN | 17 | 100.00% | 205 | 221 |
| matrix protein 1 | GFVFTLTVPSE | 11 | 100.00% | 61 | 71 |
| matrix protein 1 | TYVLSIIPSGPLKAEIAQRL | 20 | 100.00% | 9 | 28 |
| hemagglutinin | WTYNAELLVLLENERTLD | 18 | 100.00% | 436 | 453 |
| neuraminidase | FSFKYGNGVWIGRTKSI | 17 | 100.00% | 349 | 365 |
| neuraminidase | FVIREPFISCSPLECRT | 17 | 100.00% | 115 | 131 |
| matrix protein 1 | KKLKREITFHGAK | 13 | 100.00% | 101 | 113 |

| | | | | | |
|---|---|---|---|---|---|
| hemagglutinin | ELLVLLENERTLDYHDSNVK | 20 | 100.00% | 441 | 460 |
| Nucleoprotein | NPAHKSQLVWMACHSAAFEI | 20 | 95.00% | | |
| hemagglutinin | LDIWTYNAELLVLLENERTL | 20 | 95.00% | 433 | 452 |
| neuraminidase | CFWVELIRGRPKENTIW | 17 | 94.12% | 421 | 437 |
| neuraminidase | DNGAVAVLKYNGIITDT | 17 | 94.12% | 199 | 215 |
| neuraminidase | IITDTIKSWRNNILRTQ | 17 | 94.12% | 211 | 227 |
| neuraminidase | KSWRNNILRTQESECAC | 17 | 94.12% | 217 | 233 |
| neuraminidase | LDCIRPCFWVELIRGRP | 17 | 94.12% | 415 | 431 |
| neuraminidase | NQNLEYQIGYICSGIFG | 17 | 88.24% | 307 | 323 |
| neuraminidase | EKGKIVKSVEMNAPNYH | 17 | 88.24% | 259 | 275 |

**Supplemental Table 11. T cell epitopes from Cal09 in mice that were a 90 or greater percent match to bovine H5N1 conserved in Victoria, Wisconsin, or Norway H1N1 2022.** Immune epitope data was obtained by the Immune Epitope Database and Tools (IEDB) program and viral sequences were obtained from GISIAD.

| Epitope name | Epitope sequence | Epitope length | Maximum identity | Starting position | Ending position |
|---|---|---|---|---|---|
| **A/Victoria/4897/2022** | | | | | |
| nucleocapsid protein | FYIQMCTEL | 9 | **100.00%** | 39 | 47 |
| nucleocapsid protein | RLIQNSITI | 9 | **100.00%** | 55 | 63 |
| polymerase PB1 | SSYRRPVGI | 9 | **100.00%** | 703 | 711 |
| nucleocapsid protein | IQMCTELKL | 9 | **100.00%** | 41 | 49 |
| nucleocapsid protein | YSLVGIDPF | 9 | **100.00%** | 296 | 304 |
| nucleocapsid protein | RLIQNSITIERMVLS | 15 | **100.00%** | 55 | 69 |
| hemagglutinin | TYNAELLVLLENERT | 15 | **100.00%** | 437 | 451 |
| polymerase PA | ASMRRNYFTA | 10 | **100.00%** | 439 | 448 |
| polymerase PA | PSLENFRAYV | 10 | 80.00% | 224 | 233 |
| **A/Wisconsin/67/2022** | | | | | |
| nucleocapsid protein | FYIQMCTEL | 9 | **100.00%** | 39 | 47 |
| nucleocapsid protein | RLIQNSITI | 9 | **100.00%** | 55 | 63 |
| polymerase PB1 | SSYRRPVGI | 9 | **100.00%** | 703 | 711 |
| nucleocapsid protein | IQMCTELKL | 9 | **100.00%** | 41 | 49 |
| nucleocapsid protein | YSLVGIDPF | 9 | **100.00%** | 296 | 304 |
| nucleocapsid protein | RLIQNSITIERMVLS | 15 | **100.00%** | 55 | 69 |
| hemagglutinin | TYNAELLVLLENERT | 15 | **100.00%** | 437 | 451 |
| polymerase PA | ASMRRNYFTA | 10 | **100.00%** | 439 | 448 |
| polymerase PA | PSLENFRAYV | 10 | 80.00% | 224 | 233 |

| A/Norway/31694/2022 | | | | | |
|---|---|---|---|---|---|
| nucleocapsid protein | FYIQMCTEL | 9 | **100.00%** | 39 | 47 |
| nucleocapsid protein | RLIQNSITI | 9 | **100.00%** | 55 | 63 |
| polymerase PB1 | SSYRRPVGI | 9 | **100.00%** | 703 | 711 |
| nucleocapsid protein | IQMCTELKL | 9 | **100.00%** | 41 | 49 |
| nucleocapsid protein | YSLVGIDPF | 9 | **100.00%** | 296 | 304 |
| nucleocapsid protein | RLIQNSITIERMVLS | 15 | **100.00%** | 55 | 69 |
| hemagglutinin | TYNAELLVLLENERT | 15 | **100.00%** | 437 | 451 |
| polymerase PA | ASMRRNYFTA | 10 | **100.00%** | 439 | 448 |
| polymerase PA | PSLENFRAYV | 10 | 80.00% | 224 | 233 |
